# One-Pot Endonucleolytically Exponentiated Rolling Circle Amplification by CRISPR-Cas12a Affords Sensitive, Expedited Isothermal Detection of MicroRNAs

**DOI:** 10.1101/2022.05.01.490215

**Authors:** He Yan, Yunjie Wen, Song Han, Steven J. Hughes, Yong Zeng

**Affiliations:** Department of Chemistry, University of Florida, Gainesville, FL 32611; Department of Surgery, University of Florida College of Medicine, Gainesville, FL 32610; J. Crayton Pruitt Family Department of Biomedical Engineering, University of Florida, Gainesville, FL 32611; University of Florida Health Cancer Center, Gainesville, FL 32611

## Abstract

MicroRNAs (miRNAs) are a class of short non-coding RNAs that play essential roles in gene expression regulation. While miRNAs offer a promising source for developing potent cancer biomarkers, the progress towards clinical utilities remains largely limited, due in part to the long-standing challenge in sensitive, specific, and robust detection of miRNAs in human biofluids. Emerging next-generation molecular technologies, such as the CRISPR-based methods, promise to transform nucleic acid testing. The prevailing strategy used in existing CRISPR-based methods is to hyphenate two separate reactions for pre-amplification, *e*.*g*., rolling circle amplification (RCA), and amplicon detection by Cas12a/13a *trans*-cleavage in tandem. Thus, existing CRISPR-based miRNA assays require multiple manual steps and lack the analytical performance of the gold standard, RT-qPCR. Radically deviating from the existing strategies, we developed a one-step, one-pot isothermal miRNA assay termed “Endonucleolytically eXponenTiated Rolling circle Amplification with the dual-functional CRISPR-Cas12a” (EXTRA-CRISPR) to afford RT-PCR-like performance for miRNA detection. We demonstrated the superior analytical performance of our EXTRA-CRISPR assay to detect miRNAs (miR-21, miR-196a, miR-451a, and miR-1246) in plasma extracellular vesicles, which allowed us to define a potent EV miRNA signature for detection of pancreatic cancer. The analytical and diagnostic performance of our one-pot assay were shown to be comparable with that of the commercial RT-qPCR assays, while greatly simplifying and expediting the analysis workflow. Therefore, we envision that our technology provides a promising tool to advance miRNA analysis and clinical marker development for liquid biopsy-based cancer diagnosis and prognosis.

## Introduction

MicroRNAs (miRNAs) are endogenous and short non-coding single-stranded RNAs (18-23 nucleotides) that are involved in the post-transcriptional repression of messenger RNAs (mRNAs). Because they participate in various biological processes such as cell proliferation, differentiation, and cell death, dysregulated miRNAs are closely linked to the pathogenesis of diseases such as cancers.^1-4^ miRNAs were originally studied in tissues, but they have also been discovered in the blood, urine, and other body fluids, either associated with ribonucleoprotein complexes or argonaute-2, or encapsulated in exosomes,^5^ making probing circulating miRNAs a promising strategy in liquid biopsy-based cancer diagnosis, prognosis, and monitoring.^2, 5, 6^ Despite the promise of miRNAs, moving from proof-of-concept to clinical practice remains a work in progress.

One roadblock is the challenge in high-performance detection of miRNAs in biospecimens due to their short length, high sequence similarity within miRNA families, enormous concentration range in different cell types and biofluids, and complexity of associated origins and carriers.^6-9^ Therefore, there is a pressing need of ultrasensitive, specific, and robust bioassays and sensors to facilitate the development of clinically viable miRNA biomarkers of diseases.

Reverse transcription-quantitative polymerase chain reaction (RT-qPCR) has been the gold standard tool for miRNA detection. Distinct from long RNA species, such as mRNAs, short miRNAs necessitates a special RT process to incorporate extended sequences that facilitate PCR amplification and detection.^10, 11^ Stem-loop and polyadenylation (poly-A) RT assays are two commonly used approaches and broadly adapted in many commercial miRNA RT-qPCR kits.^8, 10^ Alternative to thermal cycling-based qPCR that requires sophisticated analytical procedures and instruments, many isothermal assays have been developed to advance miRNA detection,^12^ including rolling circle amplification (RCA),^13-15^ exponential amplification reaction (EXPAR),^16, 17^ loop-mediated isothermal amplification (LAMP),^18^ hybridization chain reaction (HCR),^19, 20^ and catalytic hairpin assembly (CHA).^21, 22^ Despite their merits in simplicity and even instrument-free operation, these assays have drawbacks that limit widespread clinical application, such as non-specific amplification and high background of EXPAR and LAMP, and relatively slow kinetics and low sensitivity of HCR and CHA.

Recently, CRISPR (clustered regularly interspaced short palindromic repeats) technologies have emerged as a versatile platform for developing the next-generation bioassays that combine the analytical performance of PCR and the ease of isothermal amplifications. The CRISPR-Cas12a/-Cas13a systems confer highly specific target recognition via binding with the Cas enzyme-crRNA complex and enzymatic signal amplification due to collateral cleavage (*trans*-activity) of a fluorogenic probe by the Cas enzyme activated upon target binding.^23, 24^ A variety of CRISPR assays have been reported for DNA and viral RNA detection which normally require an additional PCR or isothermal pre-amplification step to achieve desirable detection sensitivity.^25-28^ Following the same strategy, sensitive CRISPR-based miRNA assays were also developed by incorporating pre-amplification of miRNA targets by various isothermal reactions, including RCA,^29-31^ LAMP,^32^ and cascade amplification.^33^ However, these methods involving two separate pre-amplification and CRISPR-mediated readout steps require multi-step manual operations, which not only leads to complicated assay workflow and extended turnaround time, but also increases the risk of analytical variations and false results due to human error, enzymatic degradation, and cross-contaminations. It was recently demonstrated that it is possible to combine isothermal amplification and CRISPR detection in a one-pot reaction via delicately engineering the primer designs and reaction conditions.^27, 34-36^ However, it remains unclear if such strategy can be adapted to develop one-step, one-pot CRISPR assays for miRNA sensing. Alternatively, CRISPR-mediated target recognition can be integrated with other signal transduction modalities, such as electrochemical,^37^ plasmonic,^38^ and graphene field-effect transistor (gFET) sensors,^39^ enabling amplification-free nucleic acid detection. However, these approaches are not truly comparable to RT-qPCR, due to either low sensitivity without pre-amplification^37^ or the limitations arising from highly specialized devices and instruments needed. Lastly, it is worth noting that most, if not all of the existing CRISPR-based methods leverage on the *trans*-cleavage activity of Cas proteins, while the exploration of the specific *cis*-cleavage activity-the main mechanism of CRISPR-Cas systems for gene editing-for biosensing remains largely untapped.

Herein, we report a one-step, one-pot isothermal CRISPR-Cas12a assay termed “Endonucleolytically eXponenTiated Rolling circle Amplification with CRISPR-Cas12a” (EXTRA-CRISPR) for rapid, specific detection of miRNAs with RT-PCR-comparable sensitivity (**Fig. 1**). The EXTRA-CRISPR assay offers three major distinctions from the existing CRISPR-based biosensing methods. First, it presents the first strategy to simultaneously harness both *cis*-cleavage and *trans*-cleavage activities of the CRISPR-Cas12a system. Specifically, it exploits the specific *cis*-cleavage activity to transform conventional linear RCA to enable exponential amplification of target sequences, in addition to the *trans*-cleavage reaction for amplicon detection and signal amplification; Second, by engineering a modular padlock probe design and the reaction kinetics, we incorporate multiple reactions for target-mediated ligation, RCA, Cas12a binding, and nucleolytic cleavage into one collaboratively coupled reaction network, creating a robust one-step, single-tube isothermal assay that vastly simplifies the workflow for miRNA analysis; Third, this one-pot isothermal miRNA assay affords superior analytical performance that is comparable with standard RT-qPCR, including high sensitivity with a single-digit fM detection limit, single-nucleotide specificity, and rapid and flexible turnaround (from 20 min to 3 h for the entire analysis depending on targets and samples).

**Figure 1.**
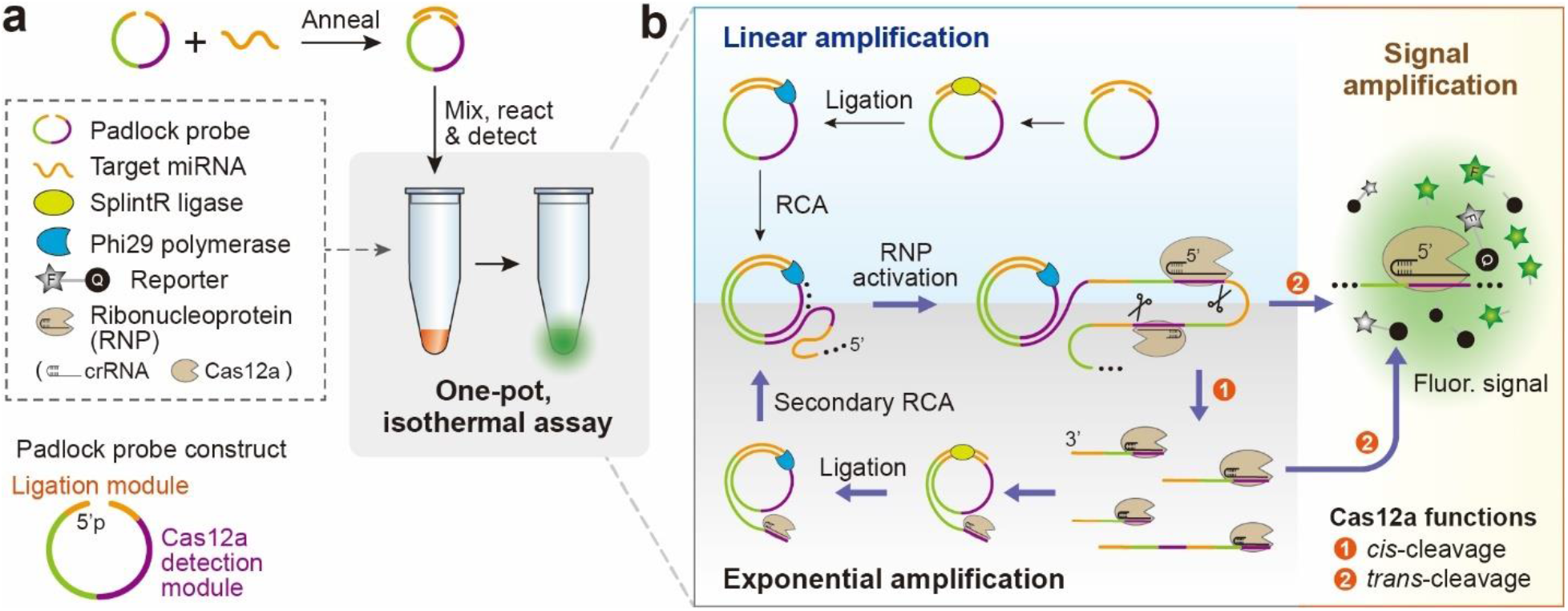
One-pot EXTRA-CRISPR miRNA assay. (**a**) The major components and workflow of EXTRA-CRISPR assay. The padlock probe for RCA is engineered with a split ligation module for target miRNA binding and a CRISPR-Cas12a detection module whose complementary sequence activates Cas12a-crRNA complex. The padlock and target sequences are annealed and added to a reaction tube containing the enzymes, reporter, and other reagents. The one-pot assay is carried out at 37 °C in a qPCR apparatus for real-time detection of the fluorescence signal. (**b**) The proposed mechanism of the EXTRA-CRISPR. This assay harnesses both *cis*- and *trans*-cleavage functions of the CRISPR-Cas12a system to convert linear RCA to an exponential amplification platform for miRNA detection. Briefly, the Cas12a RNP can bind and cleave the long ssDNA amplicon of RCA by its *cis*-activity, which generates many secondary templates containing the target sequence to initiate subsequent RCA cycles, resulting in exponential amplification of the target. Meanwhile, the amplicon-activated Cas12a RNP non-specifically cleaves the ssDNA reporters to create and amplify fluorescence signal.

As a proof-of-concept of potential applications, we adapted the EXTRA-CRISPR assay to quantifying miRNA biomarkers in extracellular vesicles (EVs) for liquid biopsy diagnosis of pancreatic ductal adenocarcinoma (PDAC). EVs, including exosomes of 50-150 nm in size, are emerging as a promising candidate for liquid biopsy because they selectively sort and transport cellular cargoes, such as proteins and nucleic acids, that mirror the physiological and pathological states of parental cells.^40-45^ EVs are considered as a major carrier of miRNAs in human biofluids and tumor-derived EVs offer a promising route to explore disease-specific miRNA signatures.^40-47^

Using the EXTRA-CRISPR, we demonstrated highly sensitive and specific profiling of a panel of four miRNA markers (miR-21, miR-196a, miR-451a, and miR-1246) in EVs derived from cell lines and clinical plasma specimens. Based on the individual EV-miRNA tests, an EV signature (EV-Sig) was devised with a machine learning method to improve the diagnostic performance for pancreatic cancer. The analytical and diagnostic performance of the EXTRA-CRSPR tests were rigorously validated by parallel RT-qPCR analysis of the same clinical samples. These results demonstrate our technology as a useful tool to advance miRNA detection and clinical development of miRNA biomarkers for liquid biopsy-based cancer diagnosis and prognosis.

## Results

### Mechanistic studies of the EXTRA-CRISPR assay

Our assay is designed to be a tri-enzymatic cascade that exploits the unique nucleolytic cleavage activities of the CRISPR-Cas12a system to create a new exponential isothermal amplification mechanism based on the robust linear RCA assay. As illustrated in **Fig. 1a**, the assay starts with the hybridization of a padlock probe with the miRNA target. The padlock probe is a 5’-phosphorylated single-strand DNA engineered to encompass two modular sequences: a ligation zone bridging the 5’- and 3’-termini with the complementary sequence to target miRNA, and a detection zone whose complementary sequence can be recognized by Cas12a-crRNA ribonucleoprotein complexes (RNP). Upon mixing with all other assay reagents in a tube, the target-splinted padlock probe will be ligated with the SplintR ligase to form a circular template for isothermal RCA reaction. Driven by phi29 DNA polymerase, RCA continuously extends target miRNA to a long linear concatemer with repeatedly complementary copies of the padlock. The Cas12a RNP pre-formed in the solution will bind to the detection zones on the long concatemer, which activates the *trans*-cleavage activity of Cas12a enzyme to non-specifically cut the fluorophore quencher (FQ)-labeled single-stranded DNA (ssDNA) reporters to produce fluorescence signal (**Fig. 1b**). At the meantime, the activated RNP can also cut the linear RCA product into short fragments via its *cis*-cleavage function. These fragments contain single or multiple complementary copies of the padlock and thus can serve as new templates to trigger many secondary RCA reactions. Such collaboratively coupled linear DNA polymerization and Cas12a *cis*-cleavage can be repeated continuously to generate the chain reactions converting conventional linear RCA to an exponential amplification assay (**Fig. 1b**). The post-cleavage RNP complexes may also cause collateral cut of the ssDNA reporters and thus further promote CRISPR signal amplification to enhance the detection sensitivity.

The development and mechanistic studies of EXTRA-CRISPR were conducted using miR-21 as the model target, which has been found overexpressed in various human tumors.^48^ The key module in our padlock probe design, the CRISPR detection zone, was verified by detecting its complementary strand with a CRISPR-Cas12a only assay. A limit of detection (LOD) at the one picomolar (pM) level was obtained (Supplementary **Fig. S1**), in line with the reported performance for preamplification-free CRISPR-Cas detection.^26, 37^ Our EXTRA-CRISPR assay is expected to produce both single-stranded RCA amplicon and the amplicon-padlock duplex both containing the crRNA-complementary sequences. RNA-guided binding with a dsDNA activator requires a protospacer-adjacent motif (PAM) to activate Cas12a for both specific *cis*-cleavage and effective non-specific ssDNA *trans*-cleavage, whereas a ssDNA does not need the PAM but yields less catalytic activity for *trans*-ssDNA cutting.^26^ Hence, we first examined the effects of CRISPR-Cas12a in our one-pot assay by tuning its activities with the PAM sequence in the padlock. As shown in **Fig. 2a**, the PAM-free Padlock-1 resulted in a significantly higher reaction rate and signal level as compared to the Padlock-2 with a PAM sequence, indicating an inhibitory effect of the PAM in the padlock for our one-pot assay. This is likely attributed to the PAM-mediated RNP binding with the amplicon-padlock duplexes that activates both *cis*-cutting of the circular templates for RCA and rapid non-specific degradation of all ssDNA species including the cleaved amplicons for the secondary RCA,^26, 34, 49^ thereby largely reducing the overall amplification efficiency. This result implies the importance of establishing proper equilibrium among the RCA and CRISPR-Cas12a cleavage reactions to catalyze efficient target amplification, as further examined below, for which a PAM-free padlock is preferred.

**Figure 2.**
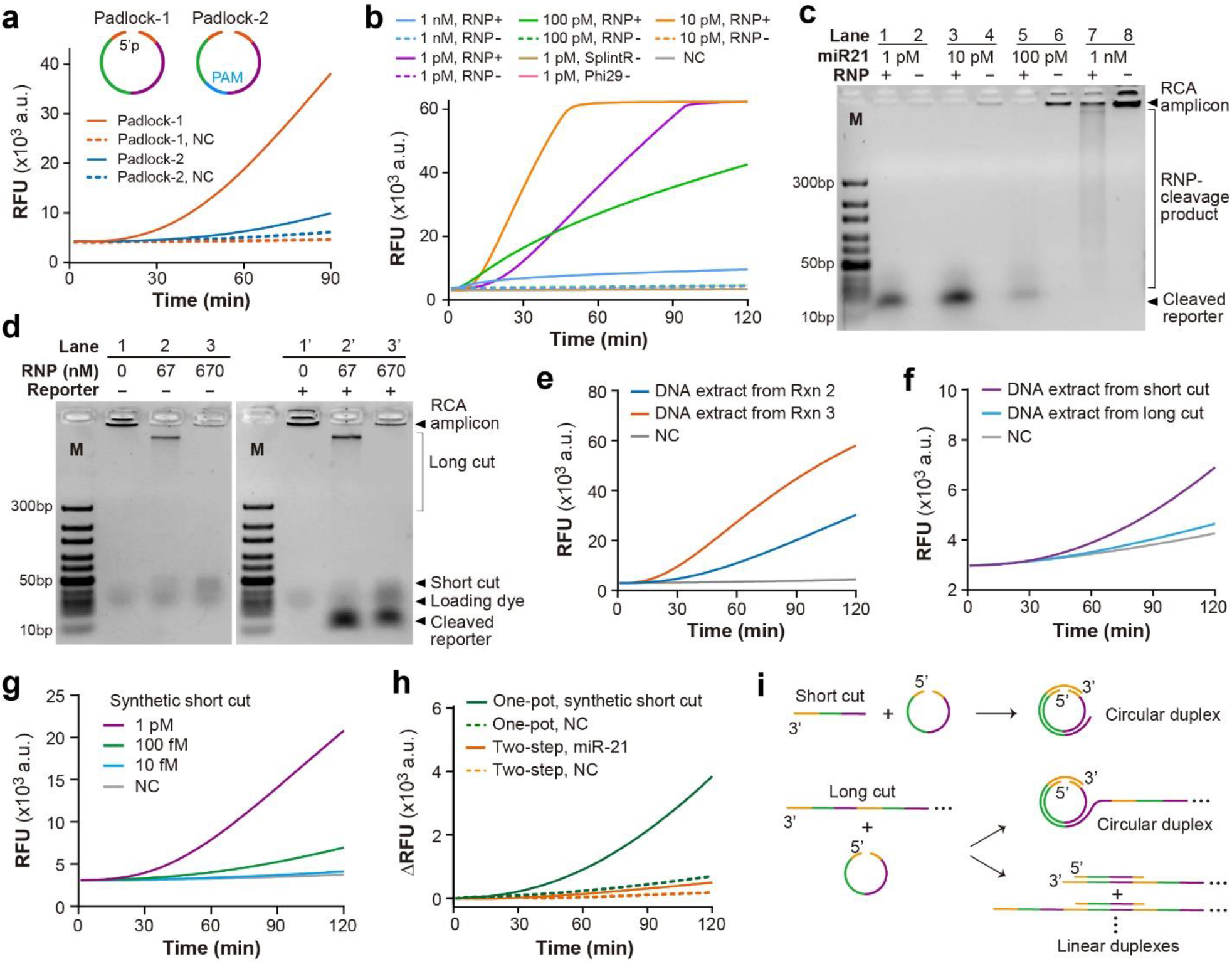
Mechanistic studies of the EXTRA-CRISPR. (**a**) Effect of the padlocks without and with a PAM sequence on miR-21 detection. The assays were conducted with 1 pM miR-21 and 100 nM of each padlock probe. NC: Negative control assay with a buffer blank. (**b**) Real-time one-pot detection of serial 10-fold dilutions of miR-21 from 1 pM to 1 nM with 1 nM Cas12a RNP. Control assays were conducted with one of three enzymes left out each time. **(c)** Gel analysis of the products from the reactions in (**b**). (**d**) Assessment of Cas12a activities on RCA amplicons in a two-step fashion. In this case, ligation-RCA was first conducted and then the products were treated with varying amounts of RNP. M: DNA marker. (**e**) EXTRA-CRISPR assays using the cleaved RCA products from Reactions 2 and 3 in (**d**) as the targets. (**f**) EXTRA-CRISPR detection of the long-cut and short-cut extracts recovered from the gel bands in Lane 2 in (**d**). Despite its less quantity, the short-cut extract yielded a faster reaction kinetics than the long cut. (**g**) Synthetic ssDNA mimicking the short-cut RCA product effectively triggers the EXTRA-CRISPR assay to produce quantitative signals. (**h**) Exponential amplification of this synthetic template was observed in the one-pot assay, as opposed to linear amplification of miR-21 in the two-step assay. The target concentration in both cases was 100 fM. (**i**) Illustration of the length-dependent binding of the Cas12a-cleaved RCA amplicons that results in differential efficiency for the secondary ligation and RCA reactions. In contrast to the short-cut amplicon, the long cuts may hybridize with the padlock sequences in the linear forms, which terminates the ligation and exponential RCA.

To facilitate the mechanistic studies, we conducted the EXTRA-CRISPR reactions at high miR-21 concentrations (1 pM to 1 nM) to permit both real-time fluorescence detection and gel electrophoresis analysis of the reaction products. **Fig. 2b** shows that our tri-enzyme assay enormously increases the detection signal with 1 pM miR-21 compared to the Cas12a only detection (Supplementary **Fig. S1**). In the control reactions with one of three enzymes left out each time, no fluorescence signal was detected, verifying the essential role of each enzyme in the one-pot EXTRA-CRISPR system. The signal intensity was observed to rise with the miR-21 concentration increased to 10 pM, but then to largely decrease at 100 pM and 1 nM (**Fig. 2b**), indicating the dynamic coupling of competing reactions associated with Cas12a in this tri-enzyme assay. The products of these reactions were analyzed with agarose gel electrophoresis. In the absence of Cas12a RNP, a band of high molecular weight DNA was detected at the edge of the sample wells and the DNA amount increased with the miR-21 input (**Fig. 2c**), which confirms successful miR-21 amplification by ligation-assisted RCA. No cleaved reporter was detected on the gel, consistent with the negative fluorescence detection seen in **Fig. 2b**. With RNP added, the one-pot reaction at a relatively low miR-21 concentration (1-100 pM) resulted in a barely detectable band of long RCA product and a weak band of small molecular weight DNA (**Fig. 2c**, lanes 1, 3 and 5), which can be presumably attributed to relatively complete cleavage of long DNA product by the activated RNP. In addition, a clear fluorescent gel band of collaterally cleaved reporter by Cas12a was detected with the intensity being enhanced at the target concentration of 10 pM and then largely reduced at 100 pM (**Fig. 2c**, lanes 1, 3 and 5), which agrees with the real-time detection results (**Fig. 2b**).

When the miR-21 concentration was further increased to 1 nM (**Fig. 2c**, lane 7), the long RCA amplicon was clearly detected, excluding the inhibition of RCA reaction as the main factor for signal suppression observed at high target concentrations. The band of long RCA amplicon was much weaker than that for the RNP-free reaction (**Fig. 2c**, lane 8) and largely smeared, which verifies Cas12a cleavage of the RCA product. Compared to the cleaved DNA bands observed for the lower target concentrations, the broad smearing indicates much less effective cleavage by the excessive DNA amplicon produced with 1 nM miR-21. While the activated Cas12a can cause both *cis*- and *trans-*cleavage of the RCA amplicon, the observed smearing bands should be mainly from the *trans*-cleavage product, because our standard gel electrophoresis assay was not sensitive enough to detect the low-level *cis-*cleaved DNA produced with 1 nM of RNP. Moreover, the cleaved-reporter band became indiscernible when the target concentration increased to 1 nM (**Fig. 2c**). Such competitive cutting of reporter versus RCA amplicon can be attributed to the non-specific ssDNase activity of Cas12a that leads to preferential cut of the RCA amplicon when a high target input initiates extremely fast RCA reaction to produce significantly more ssDNA products than the reporter. The competing effect of Cas12a *trans*-activity was verified by conducting a simple ssDNA cleavage reaction in which a transition from the reporter-dominant to padlock-dominant cleavage was observed upon the descending reporter-to-padlock ratio (Supplementary **Fig. S2**). In our assay, the padlock concentration is only 1/10 of the reporter and thus the padlock degradation by Cas12a *trans*-cutting should have minuscule effect on the EXTRA-CRISPR reaction. Together, these results suggest that our assay can produce a favorable RCA amplicon-to-reporter ratio over a broad range of target input (up to ∼100 pM, equivalent to 10^9^ copies per 20-μL reaction), enabling quantitative *trans-*cleavage of the fluorogenic reporter for accurate miRNA detection.

To further assess the Cas12a reactivities in the EXTRA-CRISPR reaction, we conducted a two-step assay in which the ligation-assisted RCA was first performed, followed by treating the amplicon with Cas12a RNP of variable concentrations. The reactions were run with the high concentrations of miR-21 (1 nM) and RNP (up to 670 nM) to facilitate the detection of both *cis-* and *trans-*cleaved DNA products. **Fig. 2d** shows that when 67 nM RNP was used, the long DNA amplicon produced by RCA (lanes 1 and 1’) were partially cleaved to yield a smeared band of long fragments and a band of short fragments (lanes 2 and 2’) which are thereafter referred to as the long and short cuts, respectively. This observation indicates enhanced cleavage of the RCA amplicon by more RNP in comparison to the assay shown in **Fig. 2c** (lane 7). Increasing the RNP concentration to 670 nM led to apparently complete digestion of the long cut into the short cut (lanes 3 and 3’). At these high levels of RNP over ssDNA substrate, the bands of *trans*-cleaved reporter were also detected (**Fig. 2d**, lanes 2’ and 3’), which is consistent with the observations for the one-pot reactions in which low levels of RCA amplicon were produced (**Fig. 2c**, lanes 1-4). A distinct observation in the two-step reactions with high-concentration RNP was an intense band of long cut (**Fig. 2d**, lanes 2 and 2’) that was barely detectable with the low level of RNP (**Fig. 2c**, lane 7). We hypothesized that this intense long cut band is mainly produced by the *cis*-activity of RNP and thus the short cut band should also contain a considerable amount of the small fragments of *cis*-cleaved RCA amplicon. These *cis*-cleavage products contain complementary copies of the padlock and may serve as new templates for secondary RCA reactions to initiate exponential amplification of target miRNA. To test our hypothesis, we investigated the ability of CRISPR-cleaved RCA products to exponentiate the linear RCA reaction. Since the ssDNA amplicon of RCA is chemically different from miRNA, we first verified that the DNA version of miR-21 yields comparable amplification efficiency with miR-21 for the EXTRA-CRISPR reaction (Supplementary **Fig. S3**). We performed the EXTRA-CRISPR assays using the DNA extracted from the two-step reactions tested in **Fig. 2d** as the template. As seen in **Fig. 2e**, significantly higher amplification was obtained with the DNA extract that contains mostly the short cut (**Fig. 2d**, lane 3) than that containing both short and long cuts (**Fig. 2d**, lane 2).

To further examine such differential reactivity of *cis*-cleaved RCA products, we extracted DNA from the separated gel bands of the short and long cuts in the lane 2 of **Fig. 2d** and input them as the templates for the EXTRA-CRISPR assays. Despite its larger quantity as detected on gel, the long-cut extract appeared to weakly trigger the tri-enzymatic reaction, while the short cut yielded much faster reaction kinetics and higher amplification signal (**Fig. 2f**). The low molecular weight of the short cut band observed on gel suggests that the *cis*-cleaved fragments in the band roughly correspond to one monomeric unit of the RCA concatemer with a length of 61 nucleotides (Supplementary **Table S1**). Thus, we assessed a synthetic ssDNA of the unit sequence as the input for the EXTRA-CRISPR reaction. **Fig. 2g** demonstrates the quantitative titration of this synthetic short cut mimic down to a concentration of 10 fM, >100-fold lower than the LOD of the preamplification-free Cas12a assay (Supplementary **Fig. S1**). The one-pot assay with the synthetic template was seen to yield notably higher signal intensity than the two-step assay involving linear amplification of miR-21 at the same concentration (100 fM, **Fig. 2h**), indicating the high efficiency of the short cut to trigger exponential RCA. Collectively, these results should verify the major contribution of the short *cis*-cleaved amplicon to exponentiating linear RCA over the long *cis*-cleaved ones, which may be explained by their length-dependent binding with the padlock probe as depicted in **Fig. 2i**. The unit sequence only binds with the termini of a padlock to form a circular duplex that initiates the exponential amplification. In contrast, the *cis*-cleaved long fragments may cause two competing effects via: 1) circularizing the padlock with the 3’-end complementary site to initiate RCA, and 2) hybridizing the padlock probes with other binding sites to form linear duplexes, which leads to the termination of exponential reaction. A long fragment has more padlock binding sites along the strand than at the terminal and thus the probability to form linear duplexes is higher than that for forming circular probes. Therefore, the ability of long cut fragments to initiate exponential RCA can be largely suppressed (**Fig. 2f**). Overall, our findings above demonstrate dynamic coupling of the *trans-* and *cis*-cleavage activities of Cas12a in the EXTRA-CRISPR assay, which enables exponential amplification of the target.

### One-pot chemistry amplifies the performance of stepwise combined reactions

To directly assess the impact of dual-activity CRISPR on the tri-enzyme reaction network, we compared miR-21 detection using the ligation-assisted RCA, tandem RCA and CRISPR, and one-pot EXTRA-CRISPR assays, under otherwise the same reaction conditions (**Supplementary Methods**). The conventional assay composed of two sequential reactions of ligation and RCA affords a high LOD at the ∼100 pM level for miR-21 (**Fig. 3a**). Tandem combination of RCA amplification with the specific and powerful Cas12a-based signal amplification vastly improved the sensing sensitivity to detect miR-21 below 100 fM (**Fig. 3b**), which is in line with the performance reported with a similar three-step RCA-Cas12a assay.^31^ Based on this three-step tandem assay, we tested a simplified version that combines ligation and RCA in one pre-amplification reaction followed by the Cas12a-powered fluorogenic readout. While simplifying the workflow, this two-step RCA-CRISPR-Cas12 approach appeared to slightly compromise the detection sensitivity and speed as indicated by the lower signal detected over a longer reaction time (**Fig. 3c**). This can be due to that the reaction conditions optimized for one enzyme may be suboptimal for others and thus the overall system. In contrast, our one-pot assay was able to overcome this challenge and confer sensitive detection of as low as 10 fM miR-21 using a protocol yet to be optimized, which notably outperforms the three-step assay based on linear RCA (**Fig. 3d**). These comparative studies support the collaborative coupling of *trans-* and *cis*-activities of CRISPR-Cas12a in our assay to drive exponential RCA and fluorogenic signal amplification simultaneously. The dynamics of the coupled tri-enzyme reaction can be affected by several major factors, which were systematically optimized as detailed below.

**Figure 3.**
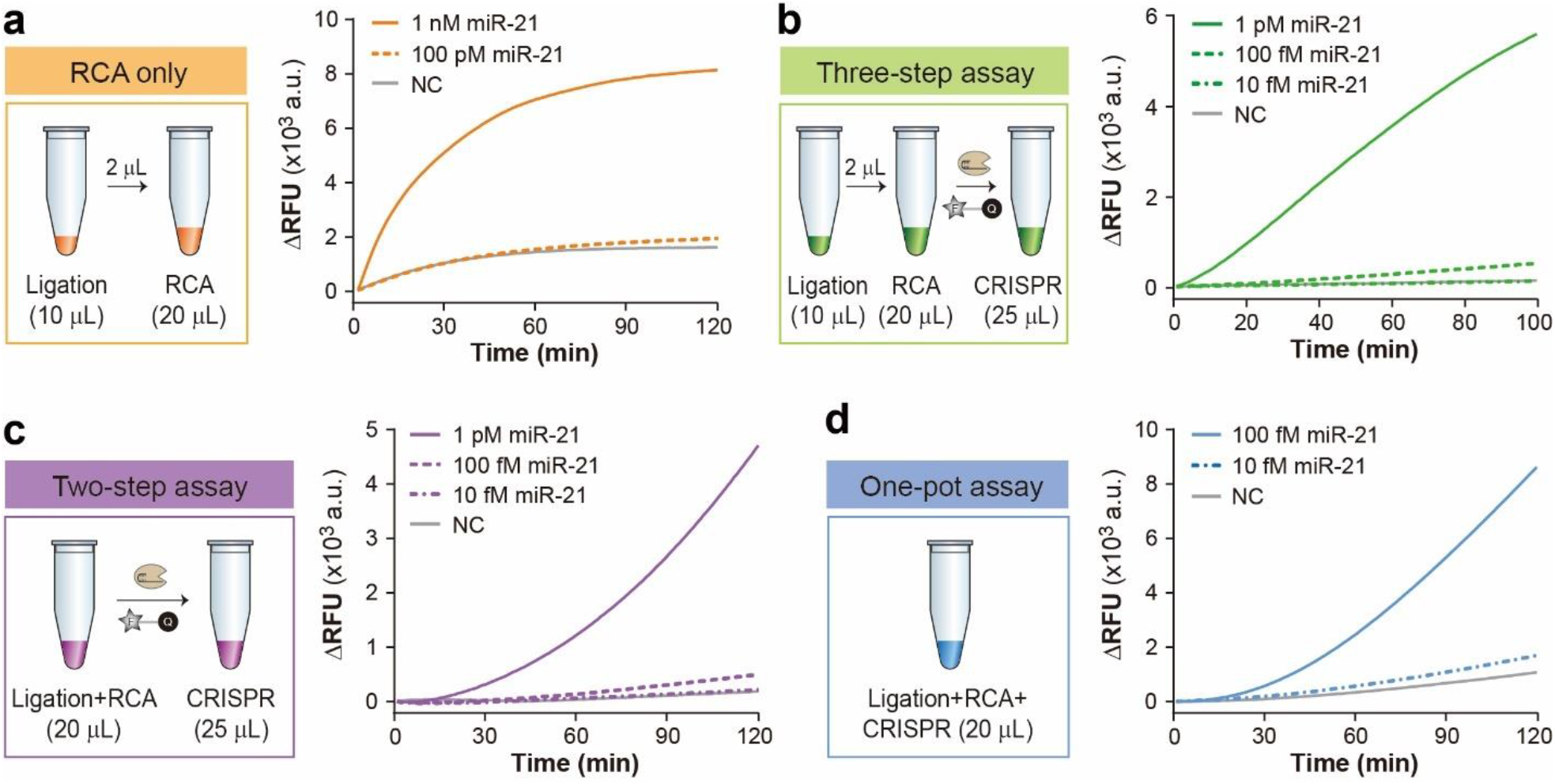
Comparison of the kinetics and detection sensitivity of RCA, one-pot EXTRA-CRISPR, and multi-step CRISPR-assisted RCA assays. (**a**) The RCA-only method involves two sequential reactions of ligation and linear RCA and can only detect 100 pM miR-21. Ligation condition: 100 nM padlock probe and 0.625 U/ μL SplintR ligase. RCA condition: 2 μL ligation product, 0.2 U/μL phi29, and SYBY Green II for detection. (**b**) The assay that connects three tandem steps of ligation, RCA, and Cas12a readout yielded a LOD of ∼100 fM. The conditions for ligation and RCA were the same as in (**a**). After RCA and denaturing of enzymes, 40 nM RNP and 0.8 μM reporter were added into the reaction. (**c**) In the two-step method, ligation and RCA were combined together, followed by fluorogenic detection using Cas12a RNP, conferring similar sensitivity with the three-step assay in (**b**). (**d**) The one-pot EXTRA-CRISPR method improves the sensitivity to detect 10 fM miR-21 prior to full optimization. Assay concentrations used in (**c**) and (**d**): 100 nM Padlock-1, 0.625 U/μL SplintR ligase, 0.1 U/μL phi29 polymerase, 1 μM reporter, and 1 nM Cas12a RNP.

### Optimization and characterization of the EXTRA-CRISPR assay

As discussed above, our padlock probe is engineered with a CRISPR detection module which guides RNP *cis*-cleavage of RCA amplicon to generate new templates for secondary RCA. Moreover, insertion of this critical module affects the overall sequence of padlock probe that is an important factor governing the efficiency of ligation and RCA reactions.^50-52^ To assess these effects and optimize the padlock probe design, we constructed three padlock probes with the CRISPR detection zone located in the right (Padlock-1), middle (Padlock-3), and left (Padlock-4) of the sequence. As displayed in **Fig. 4a**, Padlock-1 constantly generated the highest detection signal and the lowest background level over an hour of reaction, indicating its advantage to enact efficient and specific EXTRA-CRISPR reactions. Ligation of the padlock annealed to a miRNA splint is the initiating enzymatic reaction in the EXTRA-CRISPR cascade. Compared to DNA-DNA helices, RNA-splinted hybrid helices are known to be much less efficient substrates for DNA ligases, including T4 ligase that is widely used in RCA assays.^51^ To obtain an efficient ligation reaction, we compared T4 ligase and the PBCV-1 DNA ligase, also commercially branded as SplintR ligase, which was reported to provide a much higher affinity (∼300-fold KM) and turnover rate (20-fold kcat) for RNA-splinted DNA substrates.^51^ T4 ligase appeared to be ineffective to trigger the one-pot assay, whereas SplintR ligase dramatically expedited the reaction kinetics and enhanced the signal intensity (**Fig. 4b**), indicating the importance of ligation to the overall reaction kinetics and efficiency. In addition, SplintR ligase confers good stability in the assay performance over a 20-fold change of enzyme quantity (2.5 to 50 units per reaction, Supplementary **Fig. S4**).

**Figure 4.**
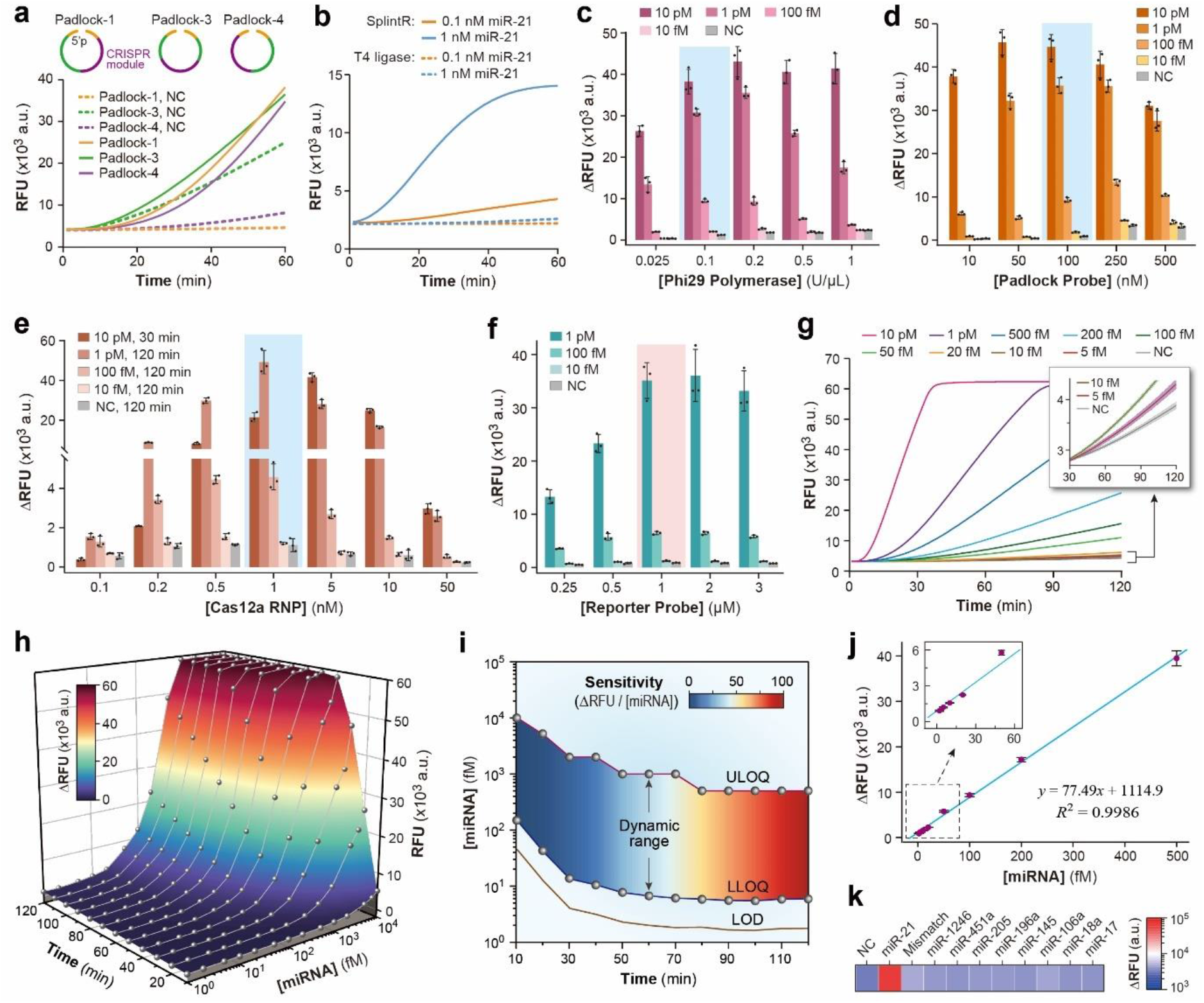
Optimization of the one-pot EXTRA-CRISPR miR-21 assay. **(a)** The position of the CRISPR module in a padlock sequence affects the detection signal and background level. The assays were conducted with 1 pM miR-21 and 100 nM padlock. **(b)** Comparison of T4 ligase and SplintR ligase for the one-pot assay. The assays were conducted with 40 U/μL T4 ligase or 2.5 U/μL SplintR ligase, 100 nM Padlock-1, 0.2 U/μL phi29 polymerase, 50 nM Cas12a RNP, and 1 μM reporter. **(c-f)** Optimization of the concentration of phi29 (**c**), padlock (**d**), Cas12a RNP (**e**), and reporter (**f**). ΔRFU, unless otherwise specified, was the signal increase from 0 min to 120 min. Error bars represent one standard deviation (S.D., *n* = 3). The selected optimal conditions were indicated by a color background. (**g**) Representative real-time curves for calibrating the one-pot assay with serial dilutions of synthetic miR-21 standards using the optimized protocol. The inset displays the curves of averaged signal for 0 (NC), 5, and 10 fM miR-21 with the shaded bands indicating one S.D. (*n* = 5). (**h**) Titration curves plotted at various time points for the assay calibration in (**g**) show a strong dependence of the assay performance on the reaction time. **(i)** Diagram of the analytical figures of merit determined by the assay calibration, including LOD, sensitivity (the slope of linear calibration curve), and linear dynamic range defined by the lower limit of quantification (LLOQ) and upper limit of quantification (ULOQ). (**j**) Linear calibration obtained with the optimal assay time of 100 min yields a LOD of 1.64 fM miR-21 calculated from 3× S.D. of the background level and a linear range from 5.47 fM to 500 fM. Error bars indicate one S.D. (*n* = 5). (**k**) Specificity of the EXTRA-CRISPR for detecting miR-21 against a single-mismatch miR-21 sequence and eight different miRNAs (1 pM each). The color intensity represents the averaged signal level of two replicates.

Next, we attempted to optimize the major RCA-related components for the EXTRA-CRISPR assay. It was found that the buffers for SplintR ligase and phi29 polymerase are more compatible with the tri-enzyme reaction and the SplintR buffer greatly outperforms the phi29 buffer at various target concentrations (Supplementary **Fig. S5**). We then optimized the phi29 polymerase concentration which is a key factor to achieving the balanced RCA and CRISPR-Cas12a cleavage reactions to catalyze exponential amplification, as reasoned before. Similar to the effect of miRNA input (**Fig. 2c**), increasing phi29 polymerase quantity will promote RCA reaction to generate higher detection signal; but excessive RCA amplicon can suppress *trans*-ssDNA cutting of reporter by RNP, resulting in reducing signal intensity. The presence of an optimal polymerase concentration was experimentally observed, which displayed a shift to the higher level for lower miRNA input (**Fig. 4c**). Similar peaking behavior was observed for the padlock probe as well because higher padlock concentration can enhance the efficiency for target binding and subsequent RCA (**Fig. 4d**). Given the low abundance of miRNAs in many biological samples, including EVs, we selected the optimal concentrations of 0.1 U/μL and 100 nM for phi29 polymerase and the padlock, respectively, which yielded the highest signal against the background for the range of 10 fM to 1 pM miR-21.

Optimization of the CRISPR components in the one-pot reaction started with assessing Cas12a RNP at a typical concentration of 50 nM used in the standard cas12a assays.^26^ However, this established assay condition led to poor signals for our assay (**Fig. 4e**), because excessive RNP can cause a termination effect, *i*.*e*., *cis*-cleavage of RCA amplicon-padlock duplexes and non-specific *trans-*cutting of the padlock, reducing the efficiency of both target amplification and fluorogenic CRISPR readout (**Fig. 2**). Thus, we screened a wide range of Cas12a RNP concentration from 0.1 to 50 nM to find a condition that maximizes the desired catalytic effect against the adverse termination effect of CRISPR-Cas12a in the one-pot exponential amplification reaction. As expected, the assay signal displayed a peaking trend as a function of the RNP concentration depending on the target input and the optimal range was narrowed down to 0.5 to 1 nM for the targeted miRNA concentration range of 10 fM to 1 pM (**Fig. 4e**). We have shown that not only can indiscriminate ssDNase activity of Cas12a cut the reporter but also degrade the padlock and linear RCA amplicon to inhibit the exponential amplification. Thus, increasing the reporter concentration would kinetically improve fluorogenic signal amplification and thermodynamically enhance the amplification efficiency by mitigating the inhibitory ssDNA cutting, while excessive reporter may raise the background signal. Indeed, the one-pot assay was seen to yield an ascending signal when the reporter concentration was increased from 25 nM to 1 μM (**Fig. 4f**). This effect was most dramatic at the high miRNA input (1 pM), which is consistent with the observed competitive Cas12a *trans-*cutting of ssDNAs (**Fig. 2c** and Supplementary **Fig. S2**). With other optimized variables, we selected a combination of 1 nM RNP and 1 μM reporter to maximize the signal to noise ratio of our one-pot assay. Collectively, these optimization studies revealed the unique dynamics of our assay which corroborates synergistic coupling of the enzymatic reactions via harnessing the dual activities of CRISPR-Cas12a.

Lastly, the analytical performance of the EXTRA-CRISPR assay was systematically calibrated with serial dilutions of synthetic miR-21 using the optimized protocol. **Fig. 4g** displays the typical curves for real-time detection over a reaction time of 2 h. The one-pot assay was able to produce signals from clearly departing from the background level at 5 fM miR-21 to reaching the limit of detector saturation for 1-10 pM miR-21, covering almost the entire dynamic range of the detector we used. The titration curves of signal versus concentration were plotted at various time points, which show a strong dependence of the assay performance on the reaction time (**Fig. 4h**). To quantitatively evaluate the impact of reaction time on the assay performance, we computed the analytical figures of merit including LOD, sensitivity (the slope of linear calibration curve), and linear dynamic range defined by the lower limit of quantification (LLOQ) and upper limit of quantification (ULOQ), which are graphically presented in **Fig. 4i**. With 20-min assay time, our method conferred a LOD of 12.3 fM which vastly outperforms the three-step assay^31^ reporting a LOD of 34.7 fM with a total of 4.5 h for both RCA and Cas12a reactions (Supplementary **Table S2**). Extending the reaction time improves the LOD and shifts the linear dynamic range down until reaching a minimal LOD of 1.64 fM with a linear range from 5.47 fM to 500 fM (*R*^2^ = 0.9986, **Fig. 4j**) at 100 min. Moreover, the calibration sensitivity of our assay can also be improved over time to better discriminate the small variations in target concentration. The analytical performance of our assay was systematically validated by the parallel measurements with gold standard RT-qPCR. It was found that our one-pot assay offers comparable detection sensitivity compared to a commercial miRCURY LNA miRNA PCR kit with which a LOD of 1.57 fM was obtained following the recommended two-step, 3-h protocol (Supplementary **Fig. S6**). To assess the specificity of our method, we extended the miR-21 assay to detect a number of non-target miRNAs including a single-nucleotide mismatched miR-21 sequence and eight human miRNAs at 1 pM each. These tests yielded a signal level of lower than 2% of the miR-21 signal except for 2.3% for the single-mismatch miR-21 (**Fig. 4k**), demonstrating the excellent specificity of our miRNA assay. Such performance can be attributed to the multi-layered protection by the specific reactions for target-padlock hybridization, ligation^52^, RCA, and Cas12a activation and *cis-*cleavage^26^. Overall, these results should strongly validate the established one-pot EXTRA-CRISPR assay for rapid miRNA detection with single-digit fM sensitivity and single-base specificity.

### Quantitative profiling of EV miRNAs for pancreatic cancer diagnosis

As a proof-of-concept demonstration of potential biomedical applications, we adapted the EXTRA-CRISPR assay to detect small EV miRNAs for diagnosis of PDAC. To this end, both cell culture media and human plasma samples were used to isolate small EVs (sEVs) and extract short RNAs from the sEV preparations, followed by parallel measurements with the one-pot EXTRA-CRISPR and two-step RT-PCR assays (**Fig. 5a**). Prior studies have identified numerous EV-miRNAs associated with human PDAC, from which we selected four serum/plasma-derived EV miRNAs that are frequently reported to be dysregulated in PDAC: miR-21,^53-56^ miR-196a,^53, 55, 57-59^ miR-451a,^54, 60-62^ and miR-1246.^57, 63^ As described above for miR-21, three EXTRA-CRISPR assays were established to detect miR-196a, miR-451a, and miR-1246 with a specific padlock probe, respectively (Supplementary **Table S3**). **Fig. 5b** presents the calibration plots for detecting the three miRNAs by EXTRA-CRISPR from which the LOD was determined to be 1.35 fM (5-500 fM linear range, *R*^2^ = 0.9974) for miR-196a, 4.14 fM (5-500 fM linear range, *R*^2^ = 0.9992) for miR-451a, and 7.96 fM (20-500 fM linear range, *R*^2^ = 0.9984) for miR-1246. Consistent with the case for miR-21, these LODs were comparable with those of the standard RT-qPCR assays which were calculated to be 1.69 fM miR-196a, 0.51 fM miR-451a, and 21.1 fM miR-1246 from the calibration curves with a threshold Ct value of 35 (**Fig. 5c** and Supplementary **Fig. S7**). Our assays were also demonstrated to afford highly specific detection of the four miRNA targets with minimal cross-reactivity between individual padlock probes and three non-targets (**Fig. 5d**). Overall, these results corroborate the excellent adaptability of our EXTRA-CRISPR method to sensitive and specific detection of miRNAs with competitive performance to RT-qPCR.

**Figure 5.**
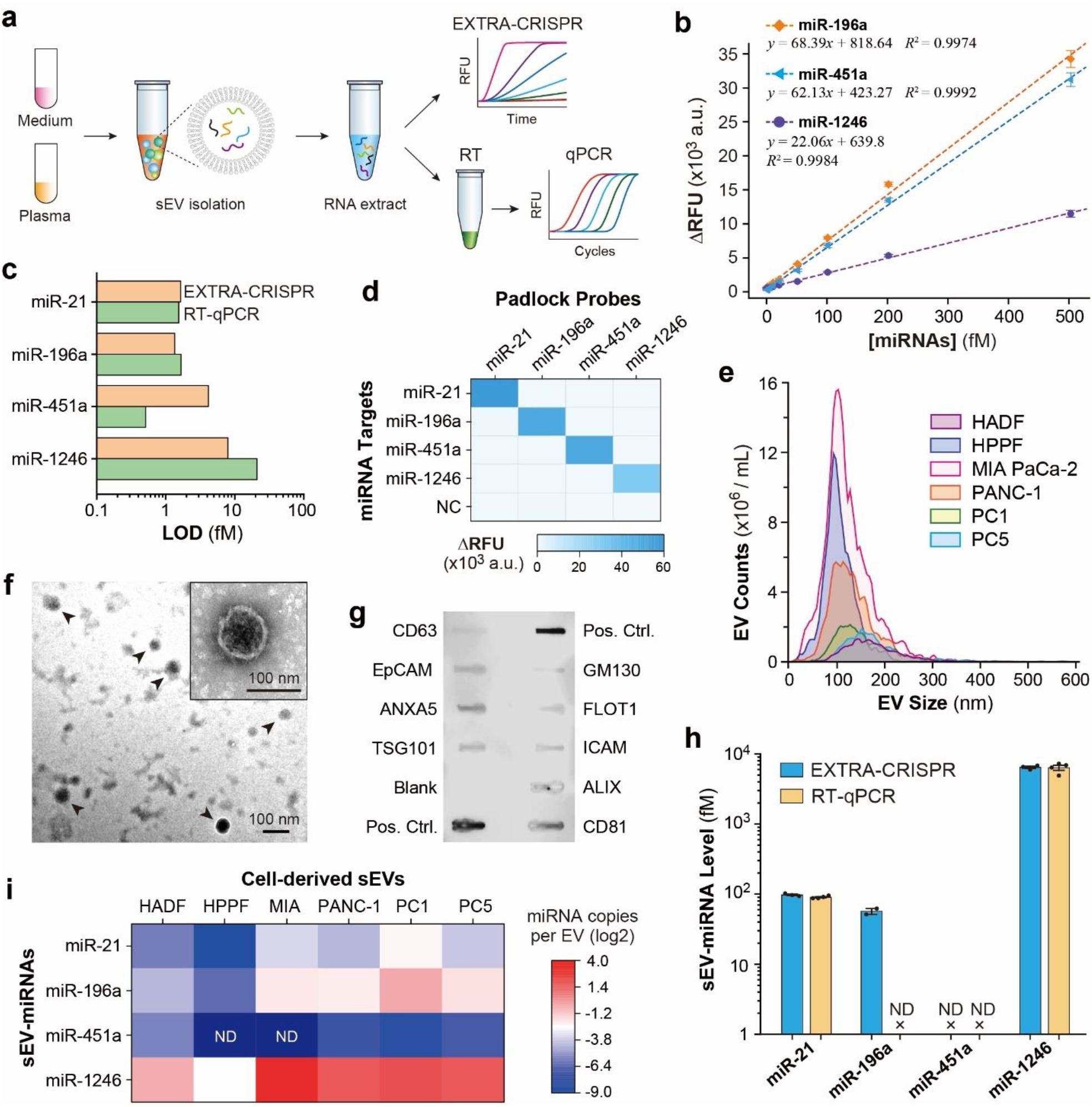
Quantitative profiling of EV-derived miRNAs. (**a**) Experimental procedure for comparative analysis of miRNAs in EVs isolated from cell culture media and human plasma, respectively, using both EXTRA-CRISPR and RT-qPCR. (**b**) Calibration curves for quantifying miRNA-196a, miR-451a, and miR-1246 by EXTRA-CRISPR. Error bars: one S.D. (*n* = 3). (**c**) Comparing the LODs for the EXTRA-CRISPR and RT-qPCR analyses of four miRNA targets. (**d**) Validation of specific analysis across four miRNAs (1 pM each) by the EXTRA-CRISPR. The color intensity denotes the averaged signal level of three technical replicates for each target. (**e**) Abundance and size distribution analyses EVs isolated from serum-free media of six control and PDAC cell lines by NTA. Normal controls: Human adult dermal fibroblasts (HADF) and human primary pancreatic fibroblast (HPPF); PDAC cell lines: MIA-PaCa-2, PANC-1, PC1, and PC5. (**f**) Representative TEM images of PANC-1 cell-derived sEVs isolated by ultracentrifugation. Inset: spherical morphology of a sEV highlighting the lipid membrane structure. (**g**) Quality assessment of isolated sEVs with a commercial antibody array. PANC-1 cell-derived sEVs were assayed to detect eight EV-associated protein markers, including CD81, CD63, FLOT1 (Flotilin-1), ICAM1 (intercellular adhesion molecule 1), ALIX (Programmed cell death 6 interacting protein), EpCAM (Epithelial cell adhesion molecule), ANXA5 (Annexin A5), TSG101 (tumor susceptibility gene 101), and a control for cellular contamination, GM130 (cis-golgi matrix protein). (**h**) Comparison of the miRNA levels of MIA-PaCa-2 sEVs determined by EXTRA-CRISPR and RT-qPCR analyses of short RNA extracted from 30 μL of purified sEVs. Error bars: one S.D. (*n* = 2 to 4 as indicated). (**i**) Heatmap of the normalized expression levels of miR-21, miR-196a, miR-451a, and miR-1246 in six cell line-derived sEVs measured by EXTRA-CRISPR. The miRNA expression level was normalized by the input number of sEV particles for each cell line. The color-coded miRNA level indicates the mean of two technical replicates of each assay. ND: not detected.

The one-pot EXTRA-CRISPR assays were assessed using purified EVs from human adult dermal fibroblasts (HADF) and human primary pancreatic fibroblast (HPPF) as the normal controls, pancreatic ductal adenocarcinoma (PDAC) cell lines (MIA-PaCa-2 and PANC-1), and PDAC patient-derived xenograft (PDX) cell lines (PC1 and PC5). Small EVs were isolated from the conditioned media by ultracentrifugation (UC)^64^ and characterized by nanoparticle tracking analysis (NTA) and transmission electron microscopy (TEM). The majority of isolated EVs displayed relatively small sizes within a range of ∼40-250 nm (**Fig. 5e**), which is typically observed for UC isolates,^65, 66^ and the morphological characteristics of sEVs (**Fig. 5f**).^65, 66^ Abundance of each cell line-derived sEVs was also measured by NTA to prepare standards for quantitative assessment of the one-pot miRNA assays. Using a commercial exosome antibody array, the quality of EV preparations was further verified by positive detection of several generic exosome protein markers (e.g., tetraspanins, Alix, ANXA5, and TSG101) and a weak signal for GM130, a control for cellular contamination (**Fig. 5g**).

We conducted miRNA profiling of MIA-PaCa-2 (MIA)-derived sEVs using the established EXTRA-CRISPR and RT-qPCR assays in parallel (**Supplementary Methods**). As depicted in **Fig. 5h**, the one-pot assay yields high consistency with RT-qPCR in measuring the concentration of miR-21 (97.42 fM by one-pot vs. 90.13 fM by RT-qPCR) and miR-1246 (6.43 pM by one-pot vs. 6.35 pM by RT-qPCR) in the samples. However, miR-196a was only detected by our one-pot assay with a determined concentration of 56.7 fM. For RT-qPCR analysis, we experienced a Ct values larger than 35 cycles for miR-196a in MIA-sEVs which prevented precise quantification. This unexpected qPCR omission can be presumably attributed to that the poly(A)-tailing method may be not robust enough to reverse transcribe miR-196a in the intricate miRNA extract where unexpected secondary structures and hybridization may hinder reverse transcription.^67^ In contrast, our one-step assay involves a denaturing and annealing procedure that eliminates potential secondary structures, empowering its robustness for detecting miRNAs in complicated biological samples. The level of miR-451a in MIA sEVs appeared to be too low to be quantifiable for both methods (**Fig. 5h**). The abundance of the targeted sEV-miRNAs for all six cell lines were measured by the one-pot assays and summarized in **Fig. 5i**. Compared to HADF and HPPF control cell lines, three miRNAs (miR-21, miR-196a, and miR-1246) were elevated in EVs from the PDAC-derived cell lines, while miR-451a was nondetectable in HPPF and MIA sEVs and very low (< ∼0.02 copies/EV) in other four cell line-derived sEVs. Relatively high miR-196a and miR-1246 levels were observed in PDAC cell-derived sEVs, which is consistent with the previous study.^57^ These findings on cell lines warrant further investigation of these sEV-miRNA markers using clinical human specimen for potential application to liquid biopsy-based diagnosis of PDAC.

We next assessed the EXTRA-CRISPR assay for measuring clinical plasma samples from cancer-free controls (*n* = 15) and patients with PDAC (*n* = 20) (Supplementary **Table S4**). As illustrated in **Fig. 5a**, the analysis workflow starts with isolating total circulating EVs from these plasma fluids (0.2 mL each) for subsequence small RNA extraction. To this end, a commercial PEG precipitation kit was used to afford faster and more efficient EV isolation than UC and hence to maximize the miRNA yield.^68, 69^ NTA analysis revealed the significant subject-to-subject heterogeneity in EV abundance varying from 6 × 10^10^ to 58 × 10^10^ EVs mL^-1^ and in mean diameters of ∼98–143 nm with the major size ranges from ∼40 to 250 nm (**Fig. 6a** and Supplementary **Fig. S8**). The isolated plasma EVs were also checked with TEM, which observed the consistent vesicle sizes and the characteristic cup or round-shaped morphologies (**Fig. 6b**). Statistical comparison showed no significant difference between the control and patient groups in the NTA measured EV levels (*P* = 0.79) and mean sizes (*P* = 0.70), respectively (Supplementary **Fig. S8**). We further assessed the EV quality with the exosome antibody array, which verified highly enriched exosomal vesicles against other cellular contaminations in these plasma EV preparations (**Fig. 6c**).

**Figure 6.**
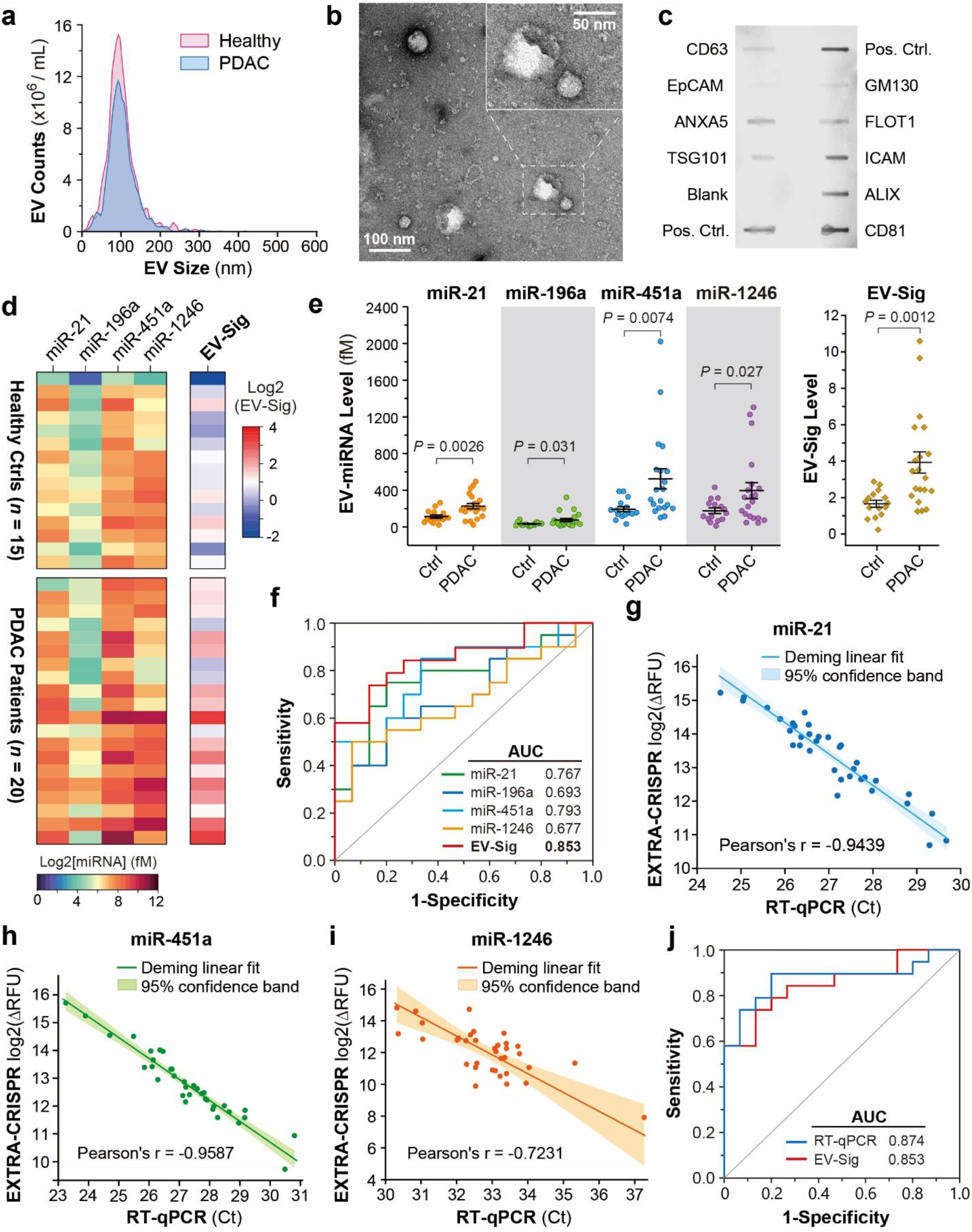
One-pot miRNA analysis for diagnosis of pancreatic cancer. (**a**) NTA of total EVs isolated by a precipitation kit from the plasma fluids of a healthy donor and a PDAC patient. (**b**) Representative TEM images of plasma sEVs from a PDAC patient. (**c**) Quality check of a patient-derived EV sample with the exosome antibody array. (**d**) Heatmap of the expression levels of individual miRNAs in the isolated plasma EVs from the PDAC patients (*n* = 20) and healthy donors (n = 15) measured by EXTRA-CRISPR. Each miRNA in each sample was tested in two technical replicates and the background-subtracted signals were adjusted by that of the positive control. The EV-Sig was defined by the weighted linear combination of four miRNAs using Lasso regression. (**e**) Scatter plots of individual sEV-miRNA markers and EV-Sig for discriminating the PDAC group from the control group. The middle line and error bar represent the mean and one s.e.m., respectively. *P* values were calculated by two-tailed Student’s *t*-test with Welch correction. (**f**) ROC curves and AUC analysis of individual sEV-miRNAs and the EV-Sig for PDAC diagnosis. (**g**-**i**) Correlation between the parallel measurements by EXTRA-CRISPR and RT-qPCR for miR-21 (**g**), miR-451a (**h**), and miR-1246 (**i**). The data points represent the mean of two replicates of each measurement by each method. Linear Deming fitting of the data points was performed to generate the linear correlation curves. (**j**) Comparing the EXTRA-CRISPR-based EV-Sig and the RT-qPCR tests of the four-miRNA panel for PDAC diagnosis. The RT-qPCR results of miR-21, miR-451a, and miR-1246 were assessed by the Lasso regression and ROC analysis. All statistical analyses were performed at 95% confidence level.

Next, we extracted and measured EV miRNAs with the EXTRA-CRISPR to quantify the levels of miR-21, miR-196a, miR-451a, and miR-1246 simultaneously. **Fig. 6d** summarizes the concentrations of the four individual plasma EV miRNA markers for each subject calculated from the calibration plots, showing their elevated expression in the PDAC cohort, in line with the previous studies.^57, 60, 70^. It is noted that compared to other three EV-miRNAs, miR-196a showed lower abundance in most of the tested clinical samples and less difference between the healthy and PDAC groups. Consistently, machine learning analysis of the data using the least absolute shrinkage and selection operator (Lasso) regression eliminated miR-196a and defined an EV signature (EV-Sig) of each subject as the weighted linear combination of miR-21, miR-451a, and miR-1246 (**Fig. 6d** and **Supplementary Methods**). As plotted in **Fig. 6e**, the EV-Sig improves the ability to differentiate the PDAC group against the healthy group (*P* = 0.0012, two-tailed Student’s *t*-test with Welch correction), compared to the individual markers (miR-21, *P* = 0.0026; miR-196a, *P* = 0.031; miR-451a, *P* = 0.0074; and miR-1246, *P* = 0.027). The diagnostic performance of these EV-miRNAs and EV-Sig was quantitatively evaluated by receiver operating characteristic (ROC) curve analysis (**Fig. 6f**). Single-marker detection yielded the modest area under the curve (AUC) values ranging from 0.677 [95% Confidence Interval (CI), 0.486–0.868] for miR-196a to 0.793 (95% CI, 0.635–0.951) for miR-451a. The EV-Sig panel combining miR-21, miR-451a, and miR-1246 greatly improves the diagnostic power to an AUC of 0.853 (95% CI, 0.726-0.979).

To validate the EXTRA-CRISPR measurements, the same RNA extracts were analyzed by the established RT-qPCR assays in parallel to quantify the four EV-miRNA markers (Supplementary **Fig. S9**). Strong correlation was observed between the two methods for detecting miR-21 and miR-451a, as indicated by the Pearson’s r of -0.9439 and -0.9587, respectively (**Fig. 6g, h**). The correlation obtained for miR-1246 was modest (Pearson’s r = -0.7231), which can be attributed to the ultralow levels of miR-1246 in most of the samples (Ct > 32) that leads to large variations in analysis (**Fig. 6i**). Consistent with the cell line results (**Fig. 5h**), the poly(A)-tailing RT-PCR assay was unable to detect EV-miR-196a in 40 thermal cycles for almost all the clinical samples tested, while the EXTRA-CRISPR assay successfully detected the low-abundance EV-miR-196a in these samples. Thus, the RT-qPCR results of miR-21, miR-451a, and miR-1246 were assessed by the ROC analysis, yielding the AUC values consistent with that of one-pot assays (Supplementary **Fig. S10**). The RT-qPCR signature combining three markers obtained through Lasso regression confers an AUC of 0.874 (95% CI, 0.757-0.991) for PDAC detection. This suggests that our assay confers a competitive performance with the RT-qPCR counterpart for plasma EV-based diagnostics targeting the same three-miRNA marker panel (**Fig. 6j**). Overall, these comparative assessments with clinical samples further support the superior sensitivity, specificity, and robustness of our method for miRNA analysis towards potential applications for liquid biopsy-based cancer diagnosis.

## Discussion

Among many isothermal methods developed for miRNA sensing, RCA is a proven technique with the advantages in simplicity, specificity, and robustness. To enhance the sensitivity limited by the linear amplification, several approaches have been explored to exponentiate RCA, including the target-primed branched RCA using a second primer^71^ and the nicking endonuclease-assisted exponential RCA.^13, 72^ Compared to these methods, CRISPR-Cas systems emerge as a compelling platform for miRNA detection owing to their substantial potential to promote both detection sensitive and specificity.^25, 26^ Nonetheless, like other CRISPR-based nucleic acid tests reported, the prevailing strategy used in the existing methods is tandem hyphenation of two separate reactions for miRNA pre-amplification and amplicon detection by Cas12a *trans*-cleavage (Supplementary **Table S2**). A challenge in combining RCA (and other amplification assays) and CRISPR assays may be attributed to the double-edged effects of CRISPR-Cas12a, as revealed by our mechanistic studies (**Fig. 2**), where indiscriminate *trans*-cleavage of Cas12a causes degradation of the essential ssDNA reactants (i.e., padlock probe and secondary templates) to suppress the exponential amplification. To overcome this issue, we discovered a strategy that combines engineering of the CRISPR-reactive padlock probe and balancing the reaction kinetics and equilibria to promote the desired Cas12a functions in a complex tri-enzymatic reaction network (**Fig. 2a** and **Fig. 4**). This novel strategy allowed us to establish the first one-step, one-pot isothermal assay that collaboratively couples RCA with the CRISPR-Cas12a system to enact exponential amplification and fluorogenic detection of miRNAs. Moreover, our work suggests the feasibility of harnessing both *cis-* and *trans-*activities of CRISPR-Cas systems for biosensing, paving a new way for developing the next-generation CRISPR diagnostics.

High sensitivity is essential for detecting miRNA signatures of tumors in biospecimens. The concentrations of miRNA markers in biofluids can be at the picomolar level or even lower, especially at the early disease stages.^73^ Relevant to this work, the EV population is considered as a major carrier of miRNAs in human biofluids and exosomes secreted by tumor cells offer a promising route to explore disease-specific miRNA signatures.^74-76^ However, it has been shown that the averaged load of a miRNA target in EVs can be as low as 10^−5^ copies per vesicle,^13, 77^ which makes sensitive miRNA analysis essential to clinical biomarker development. Our EXTRA-CRISPR miRNA assay offers a single-digit fM sensitivity and single-nucleotide specificity, which is comparable with RT-qPCR,^78^ while greatly simplifying and expediting the analysis workflow. Our assay is also cost-effective with the material cost estimated to be as low as $0.6 per test at the research scale. Such combination of superior analytical performance, one-pot contamination-proof operation, and low cost presents our method as a radical improvement to the existing CRISPR-based miRNA assays (Supplementary **Table S2**).^29, 31, 33, 79^

PDAC that accounts for ∼90% of all pancreatic neoplasms is an extremely aggressive and lethal gastrointestinal malignancy that is projected to be the second-leading cause of cancer-related mortality by 2030, with an overall 5-year survival rate of 11% and an incidence increase rate of 0.5%-1.0% per year .^80-82^ Due to its asymptomatic course, rapid progression, and delayed clinical presentation, PDAC is considered a silent killer, and most patients were diagnosed at an advanced stage.^83, 84^ Early detection of PDAC at resectable stages has profound impact on changing the malignancy’s poor survival figures.^85, 86^ However, no reliable screening method, either molecular or imaging-based, exists to date to allow accurate early detection of asymptomatic PDAC patients. Serum carbohydrate antigen 19-9 (Ca19-9) detection is the most extensively adapted marker for PDAC diagnosis which was approved by the U.S. Food and Drug Administration (FDA) in 2002. Ca19-9 can help with PDAC prognosis and monitoring, however, it is not recommended as a biomarker for early detection due to its low positive predictive value (0.5–0.9%).^87^ Increasing evidence has indicated the promising potential of circulating EVs as a rapid, minimally invasive, efficient, and cost-effective liquid biopsy in developing diagnostic biomarkers of PDAC.^70^ Using PDAC as the disease model, we assessed the feasibility of the EXTRA-CRISPR assay for clinical analysis of miRNA markers in circulating EVs for PDAC diagnosis. Our assay shows good compatibility with two commonly used EV isolation methods. Targeting four PDAC-related markers (miR-21, miR-196a, miR-451a, and miR-1246), we demonstrated highly sensitive and specific miRNA profiling of EVs derived from cell lines and clinical plasma specimens. While individual EV-miRNA tests only yielded modest diagnostic power, an EV signature (EV-Sig) can be defined from the four-marker panel by machine learning analysis to improve the PDAC detection with an AUC of 0.853 (**Fig. 6f**). This diagnostic performance is comparable with that of the serum Ca19-9 test previously reported,^88^ which supports the clinical potential of EV-miRNA markers for PDAC diagnosis. Rigorous validation by RT-qPCR analysis of the same samples was conducted in parallel to the one-pot assays and showed strong correlation in both analytical and diagnostic performance between the two methods, confirming the robustness of the EXTRA-CRSPR assay. It is noted that the primary goal of this proof-of-concept clinical study was to assess a new bioassay rather than the biomarkers. To this end, we simply gathered the panel of four miRNAs highly associated with PDAC from literature. An improved EV-Sig, constituted with an optimal miRNA biomarker panel for PDAC diagnosis, will no doubt enhance the diagnostic power of our new one-pot assays. Future work of large-scale screening and validation of circulating EV-miRNAs using the developed EXTRA-CRSPR assay could facilitate the identification of potent EV-miRNA biomarker candidates in PDAC, and beyond.

Overall, featuring a combination of superior analytical performance, easy operation, rapid reaction, and low cost, the EXTRA-CRISPR provides a competitive alternative to standard RT-qPCR for miRNA analysis in biological and clinical samples. The simplicity of the method would greatly facilitate future instrumentation or microfluidic miniaturization and integration to fully automate the workflow, further increase analysis throughput and reproducibility, and reduce sample consumption and turnaround time. For instance, our assay can be readily integrated with microfluidic EV isolation^89, 90^ to greatly streamline the analytical pipeline and improve the performance for clinical analysis of EV miRNA markers. In principle, our CRISPR-enabled isothermal amplification technology could provide a versatile platform for developing new biosensors for other types of biomarkers such as proteins and small molecules, using aptamers or other nucleic acids as a bridge.^91, 92^ Therefore, our method provides a useful tool to facilitate the development of next-generation biosensing technologies and new clinical biomarkers. Therefore, we envision that this new approach would open immense opportunities for developing the next-generation CRISPR diagnostics to address the needs in broad areas, including biological research, clinical lab diagnosis and POC testing.

## Methods

### Materials

RNA and DNA oligos were purchased from IDT (Integrated DNA Technologies), SplintR buffer, dNTP (10 mM), BSA (20 mg/mL), phi29 polymerase (10 U/μL), SplintR ligase (25 U/μL), cas12a (1 μM) and ATP are purchased from NEB (New England Biolabs Inc). SYBR Green II dye, SYBR gold nucleic acid gel stain, agarose, and 10× TBE buffer were purchased from Thermo Fisher Scientific. SeraMir™ exosome RNA amplification kit (RA806A-1), SeraMir exosome RNA column purification kit (RA808A-1), and Exo-Check™ exosome antibody array (EXORAY200B-4) were purchased from SBI (System Biosciences). miRCURY LNA™ RT kit, miRCURY LNA™ SYBR Green PCR kit, and the miRNA specific primers were purchased from Qiagen. Pierce™ rapid gold BCA protein assay kit (A53226) was purchased from ThermoFisher Scientific. WesternBright™ Sirius™ chemiluminescent HRP substrate was purchased from advansta.

### Procedures of EXTRA-CRISPR assay

For the preparation of RNP, 2 μL of water, 0.5 μL of 2.1 buffer (NEB), 1 μL of cas12a (1 μM) and 1.5 μL of crRNA (1 μM) was incubated together at 37°C for 30 min. The usage of each component can be adjusted according to the experimental consumption. For a 20-μL reaction system, padlock (2 μL) and miRNA (2 μL) in buffer (0.4 μL) are first denatured at 80°C for 5 min and cooled down gradually to room temperature, and then added to the reaction system containing other components (15.6 μL in total, with 0.8 μL of dNTP, 0.2 μL of BSA, phi29 polymerase, SplintR ligase, RNP, reporter DNA, 1.6 μL of 10x buffer and water). After mixing, the assay was performed in a qPCR device (Bio-Rad, CFX connect real-time system) at 37°C for a certain period, and the fluorescence intensity is monitored every 1 minute. The final concentration of miRNA in the 20-μL reaction system was used to determine the analytical sensitivity.

### Procedures of RCA only, two-step method and three-step method

In the RCA only method, padlock (1 μL), miRNA (1 μL), and SplintR buffer (0.2 μL) were first denatured at 80°C for 5 min and cooled down gradually to room temperature before being added to the ligation system which contains 0.8 μL of SplintR buffer, 0.25 μL of SplintR ligase and 6.75 μL of water. Ligation was performed at 37°C for 2 h and then 65 °C for 10 min. Then 2 μL of the ligation product was mixed with the 20-μL RCA reaction system which contains 2 μL of phi29 buffer, 0.4 μL of phi29, 0.2 μL of BSA, 0.8 μL of dNTP, 2 μL of 10x SYBR Green II, and 12.6 μL of water. RCA was performed at 37°C for 2 h in the qPCR device and monitored every 1 minute. The miRNA concentration in the ligation system was used to determine the analytical performance. In the two-step method, 2 μL of padlock, 2 μL of miRNA and 0.4 μL of SplintR buffer were first denatured and annealed and then transferred to the 20-μL ligation-RCA system which contains 1.6 μL of SplintR buffer, 0.8 μL of dNTP, 0.2 μL of BSA, 0.2 μL of phi29, 0.5 μL of SplintR and 12.3 μL of water. The reaction was conducted at 37 °C for 2 h and then 65 °C for 10 min, next, 5 μL of RNP and 0.2 μL of the reporter (100 μM) were added to the system and incubated at 37°C for 2 h in the qPCR device. The miRNA concentration in the ligation-RCA system was used to determine the analytical performance. In the three-step method, ligation and RCA step were the same with the RCA only method, after RCA, the enzyme was inactivated at 65 °C for 10 min followed by the addition of 5 μL of RNP and 0.2 μL of the reporter (100 μM), the final step was carried out at 37°C for 2 h in the qPCR device. The miRNA concentration in the ligation system was used to determine the analytical performance.

### Agarose gel electrophoresis

The gel electrophoresis experiments were performed using the 3% agarose gel in TBE buffer at 110 V for 1 h. After that, the gel was immersed into the TBE buffer containing SYBR gold dye for 30 min. The image was captured with an imager (Odyssey, LI-COR).

### RT-qPCR detection of miRNAs

The 10-μL of the reverse transcription contains 2 μL of 5× miRCURY RT reaction buffer, 1 μL of 10× miRCURY RT enzyme mix, 1 μL of miRNA, and 6 μL of water. The RT system was incubated at 42°C for 60 min and then inactivated at 95°C for 5 min. The cDNA solution was diluted by 60× before being added to the qPCR system which contains 5 μL of 2× miRCURY SYBR green master mix, 1 μL of the PCR primer, 3 μL of cDNA template, and 1 μL of water. Temperature program: 95°C for 2 min, 40 cycles of 95°C for 10 s, and 56°C for 60 s.

### Cell culture

Procedures for cell culture and the harvest of EV-contained conditioned SFM (serum-free medium) were as previously described.^93, 94^ Briefly, immortalized PDAC cell lines (MIA-PaCa-2 and PANC-1), primary human xenograft-isolated PDAC cell lines (PC1, PC5), as well as human fibroblast cell lines (HADF and HPPF) were cultured in DMEM-F12 supplemented with 10% FBS and 1% antibiotic antimycotic solution. Culture medium was replaced by DMEM-SFM when ∼85% confluence was reached. Conditioned-SFM were collected after 48 h and stored at -80°C until analysis.

### EV isolation from cell culture medium

All the centrifugations were performed at 4°C with the following procedure: 300 g for 10 min to remove cells, 2,000 g for 20 min to remove dead cells and cell devirs, 10,000 g for 30 min to remove large EV particles (e.g., microvesicles), and 100,000 g for 2 h to collect the sEV pellet. The sEVs were resuspended in 100 μL of PBS for the following NTA analysis and miRNA extraction using SeraMir exosome RNA column purification kit.

### EV isolation, miRNA extraction and detection from plasma samples

Human plasma was obtained from Clinical and Translational Science Institute, University of Florida (IRB202200150). Small EVs were isolated according to the protocol of the SeraMir™ Exosome RNA Amplification kit. First, thaw the plasma samples on ice and centrifuge at 3000 g for 15 min to remove cells and cell debris, followed by the centrifugation at 18000 g for 30 min to remove large vesicles. All the centrifugations were performed at 4°C. Then, 200 μL of the plasma was diluted to 250 μL with PBS before adding 60 μL of ExoQuick and incubated at 4°C for 30 min. After incubation, the plasma was centrifuged at 13000 rpm for 2 min to collect the pellet. EV miRNA was purified and eluted in 30 μL of water according to the manufacturer’s protocol for the following analysis. For the detection of miR-21, miR-196a, and miR-1246, 2 μL of the miRNA extract were added to the one-pot assay. As the expression level of miR-451a is extremely high, the extract was diluted 10 times before being added to the one-pot assay for the precise quantitation of miR-451. The signals using one-pot assay were subtracted by the corresponding background and normalized by miRNA positive controls. miRNA concentrations were tested by RT-qPCR simultaneously, the extract of was also diluted 10 times before being added to the reverse transcription system when detecting miR-451a. The cDNA solution was diluted 30× to perform the qPCR reaction.

### Transmission electronic microscopy (TEM)

Negative staining method is used for EV imaging. EVs were fixed in 2% paraformaldehyde (PFA) for 5 min and loaded on 200 mesh copper grids. TEM was performed at the Electron Microscopy Core of University of Florida on a Hitachi 7600 transmission electron microscope (Hitachi High-Technologies America, Schaumburg, IL) equipped with a MacroFire® monochrome progressive scan CCD camera (Optronics, Goleta, CA).

### NTA analysis

Particle number and size distribution of EVs isolated from medium and plasma samples were determined by nanoparticle tracking analysis (NTA) using a ZetaView system (Particle Metrix Inc.). Samples were diluted in PBS to an acceptable concentration, according to the manufacturer’s recommendations.

### Antibody arrays for the detection of exosomal biomarkers

The protein biomarkers of exosomes isolated from the PDAC cell line and PDAC patient’s plasma were tested using the Exo-Check™ exosome antibody array. Isolated EVs were resuspended in PBS and the amount of protein in EV samples was determined by Pierce™ rapid gold BCA protein assay kit. 50 μg protein was used for the antibody array according to the manufacture’s protocol. Briefly, the sample was lysed by lysis buffer and 1 μL of labeling reagent was added to the lysate followed by the incubation at room temperature for 30 min with constant mixing. After removing excess labeling reagent, lysates were mixed with 5 mL of blocking buffer. The blocking buffer/labeled exosomes lysate mixture was then incubated with the antibody precoated membrane at 4°C overnight on a shaker. Next day, the membrane was washed carefully by wash buffer and incubated with 5 mL of detection buffer at room temperature for 30 min. After removal of the detection buffer and washing, the membrane was developed using the WesternBright™ Sirius™ chemiluminescent HRP substrate. The image was captured using the LI-COR imager (Odyssey) with exposure time of 1 min.

### Statistical analysis

Mean, standard deviation, and standard error were calculated with standard formulas in Excel. To compare the patient and control groups, a two-tailed Student’s *t*-test with Welch correction was performed with a significance level of *P* < 0.05. ROC analyses were performed to determine the AUC values using the OriginPro software (OriginLab Corporation). Machine learning analysis of the miRNA markers was conducted by fitting the data with the least absolute shrinkage and selection operator (Lasso) paths for regularized logistic regression,^95^ in which with the tuning parameter (λ) selected by the Leave-one-out cross-validation.^96^ The estimated coefficients obtained for the markers by Lasso regression were used to create a weighted linear combination of the markers. Lasso regression was performed using the JMP Pro software (JMP Statistical Discovery LLC). A 95% confidence level was used for all statistical analyses.

## Acknowledgement

We thank the UF Clinical and Translational Science Institute (CTSI) Biorepository Facility for providing human plasma specimen. We thank Dr. Yutao Li and Gerik Tushoski for helping with TEM analysis and culturing cells and collecting conditioned medium, respectively. This study was supported in part by the grants R01 CA243445, R33 CA214333, R33CA252158A1, and R01CA260132 from National Institutes of Health.

## Supplementary Information

Additional information and results (Fig. S1-S10 and Tables S1-S4).

## Competing financial interests

H.Y. and Y.Z are co-inventors on a US provisional patent application based on this work (no. 63/313,870; title: One-pot Endonucleolytically Exponentiated Rolling Circle Amplification by CRISPR-Cas12a). Y.Z holds equity interest in Clara Biotech Inc. and serves on its scientific advisory board. All other authors declare no competing interest.

## Author contributions

Y.Z. conceived and supervised the project; H.Y. and Y.Z. designed the research; H.Y. performed technology development, mechanistic study, analytical characterization, and clinical validation; H.Y. and Y.W. performed isolation and NTA analysis of EVs; S.H. conducted cell culture, provided culture media, helped with TEM analysis, and involved in experiment design and discussion; S.J.H. assisted clinical studies; H.Y. and Y.Z. analyzed the data and wrote the manuscript. All authors edited the manuscript.

## Data availability

The authors declare that all data supporting the findings of this study are available within the paper and its Supplementary Information. The raw and analysed datasets generated during the study are available for research purpose from the corresponding author on reasonable request.

## Notes

### Competing Interest Statement

The authors have declared no competing interest.

