## Supplementary information for "One-Pot Endonucleolytically Exponentiated Rolling Circle Amplification by CRISPR-Cas12a Affords Sensitive, Expedited Isothermal Detection of MicroRNAs"

### Supplementary Figures and Tables

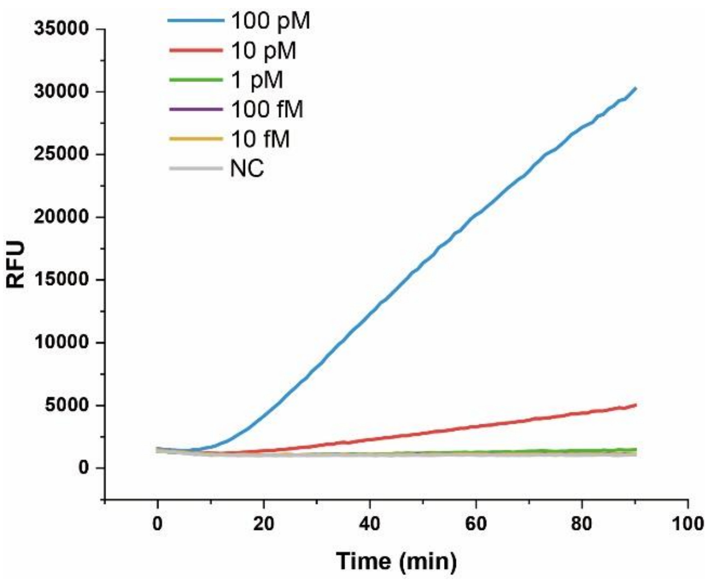


**Figure S1. ssDNA detection by CRISPR-Cas12a only.** The assay consists of a ssDNA target, Cas12a-crRNA RNP, and reporter and was conducted at 37°C. The concentrations of RNP and reporter were both 50 nM. The target containing a complementary fragment to crRNA is: GCCGGGGTGGTGCCCATCCTGGTCGAGCTGGACGGCGACGTAAACGGCCACAAG.

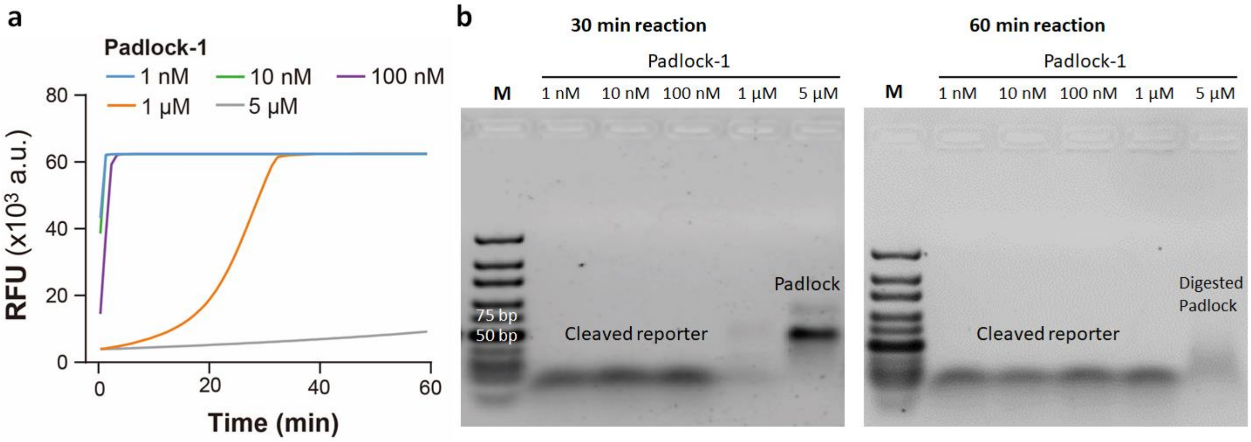


**Figure S2. Competing effects of Cas12a *trans*-cleavage activity on the padlock and reporter.** (**a**) Real-time fluorescent signals of Cas12a cleavage assays with the padlock and reporter sequences mixed at variable concentration ratios. The padlock of a concentration varied from 1 nM to 5 μM was mixed with 1 μM of the reporter and incubated with 50 nM of the activated Cas12a RNP for 1 h. (**b**) Gel analysis of the products formed at 30 and 60 mins in the reactions shown in (**a**). As the Padlock-to-reporter concentration ratio increased, both fluorescence signal (**a**) and the quantity of cleaved reporter detected on gels (**b**) were seen to decrease indicating suppression of the *trans*-cleavage of the reporter by Cas12a RNP. Meanwhile, the cleavage of the padlock was observed, which increases with reaction time, as shown in (**b**).

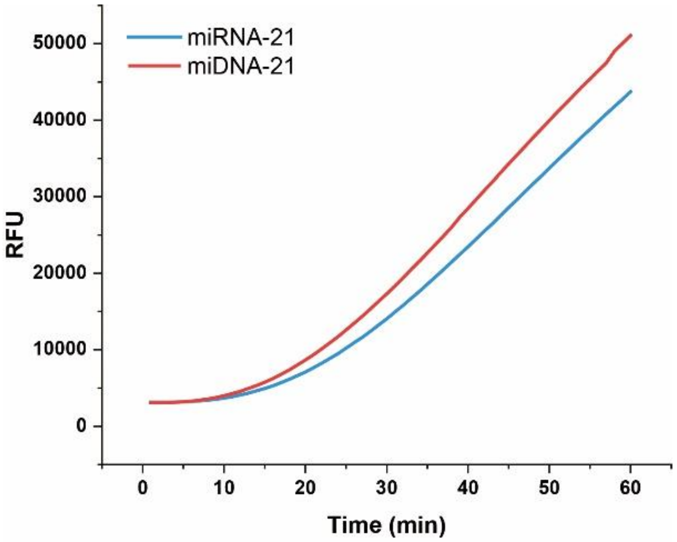


**Figure S3**. **EXTRA-CRISPR reaction using miRNA and ssDNA as the ligation templates.** Either miR-21 and its DNA version (miDNA-21) were tested to initiate the one-pot reaction. SplintR ligase can also effectively ligate the padlock probe with a DNA template. The assays were conducted in triplicate and the averaged signals were plotted. The concentration of padlock, SplintR ligase and phi29 polymerase, reporter, and RNP was 100 nM, 0.625 U/ μL, 0.1 U/ μL, 1 μM, and 1 nM, respectively.

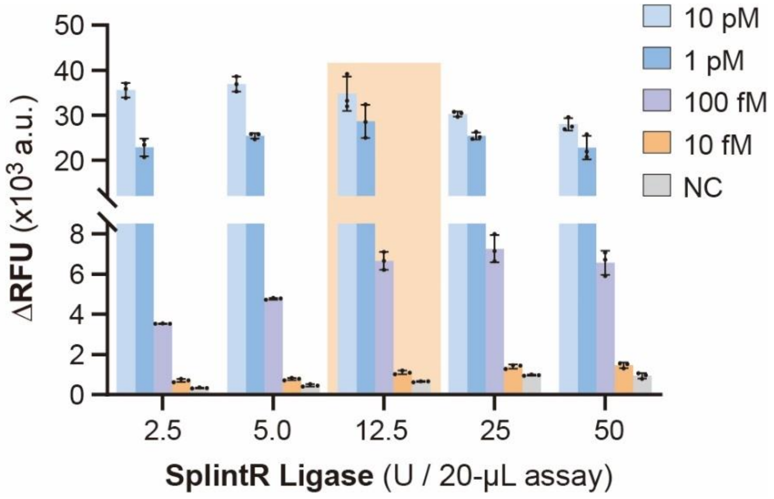


**Figure S4. Effect of SplintR ligase concentration on the one-pot assay**. SplintR ligase shows good stability in the assay performance over a 20-fold change of enzyme input from 2.5 units to 50 units per 20- μL reaction.

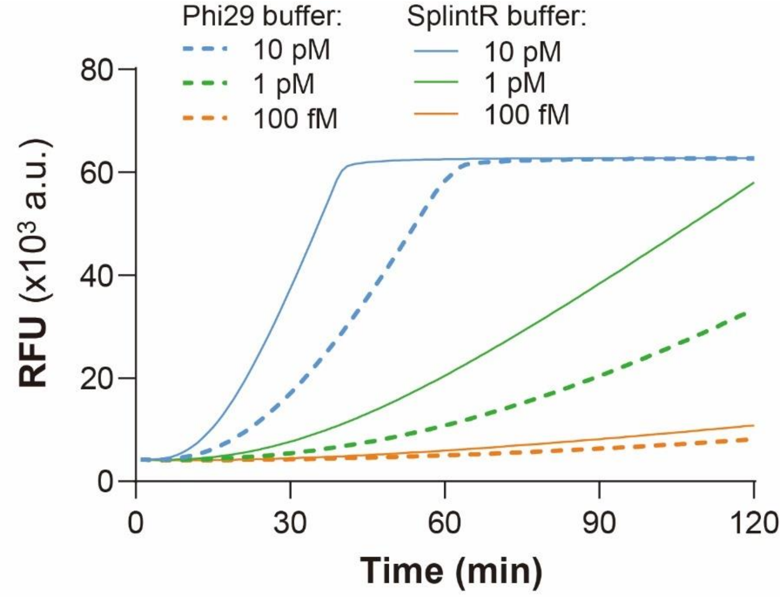


**Figure S5. Optimization of the reaction buffer for the EXTRA-CRISPR assay**. Different buffers for individual enzymes were tested for detection of serial dilutions of miR-21. The SplintR buffer appeared to largely outperform phi29 buffer, offering faster kinetics and higher detection signal. The concentration of padlock, SplintR ligase, phi29 polymerase, reporter, and RNP was 100 nM, 1.25 U/μL, 0.2 U/μL, 1 μM, and 5 nM, respectively. ATP was added to the phi29 buffer at the same concentration as in the SplintR buffer. All assays were conducted in two replicates.

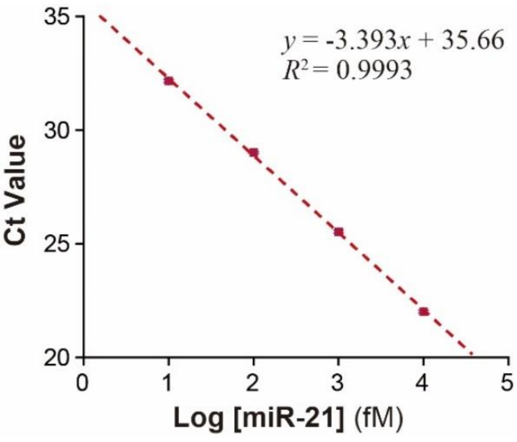


**Figure S6. Calibration of the RT-qPCR assay for miR-21 Detection.** Serial 10-fold dilutions of miR-21 were detected using the miRCURY LNA miRNA PCR kit. A LOD of 1.57 was calculated from the calibration curve with a threshold Ct value of 35. The data points and error bars indicate the mean and one S.D. (*n* = 3), respectively.

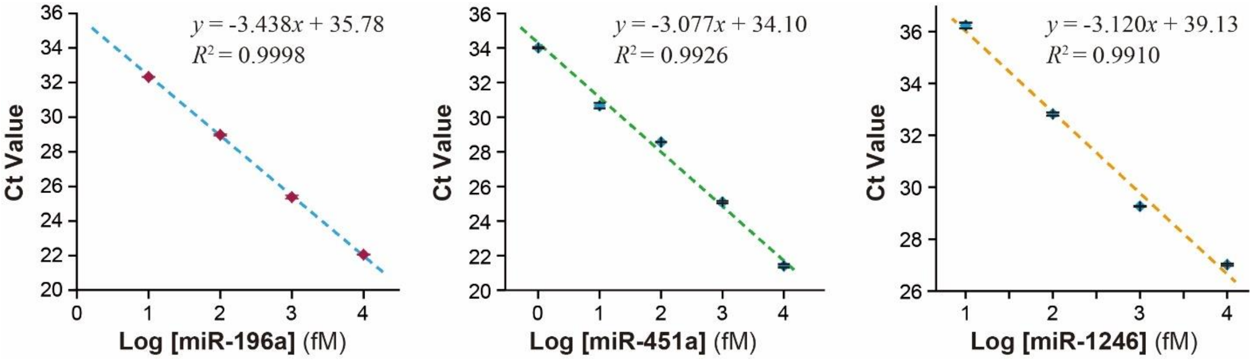


**Figure S7. Calibration curves for the RT-qPCR assays for miR-196a, miR-451a, and miR-1246.** LODs for miR-196a, miR-451a, and miR-451a were calculated to be 1.69 fM, 0.51 fM, and 21.1 fM, respectively, using the calibration curves with a threshold Ct value of 35. The data points and error bars indicate the mean and one S.D. (*n* = 2), respectively.

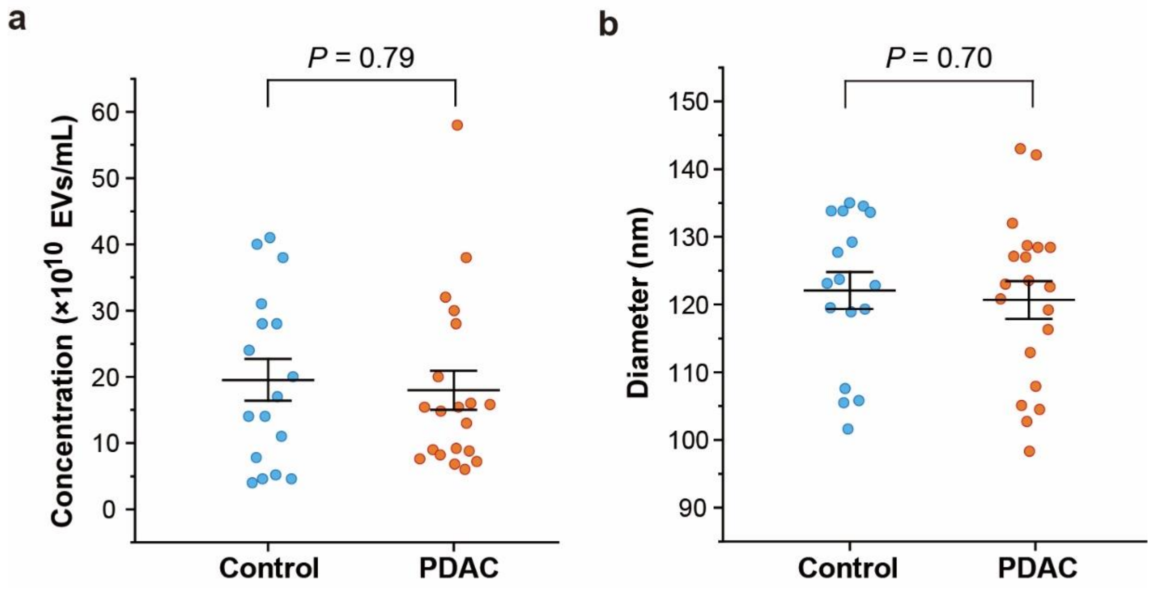


**Figure S8. Scatter dot plots for (a**) the abundance and (**b**) mean diameter of EVs purified from the control and patient plasma samples measured by NTA. *P* values were calculated by two-tailed Student's *t*-test with Welch correction. The middle line and error bar represent the mean and one s.e.m., respectively. All statistical analyses were performed at 95% confidence level.

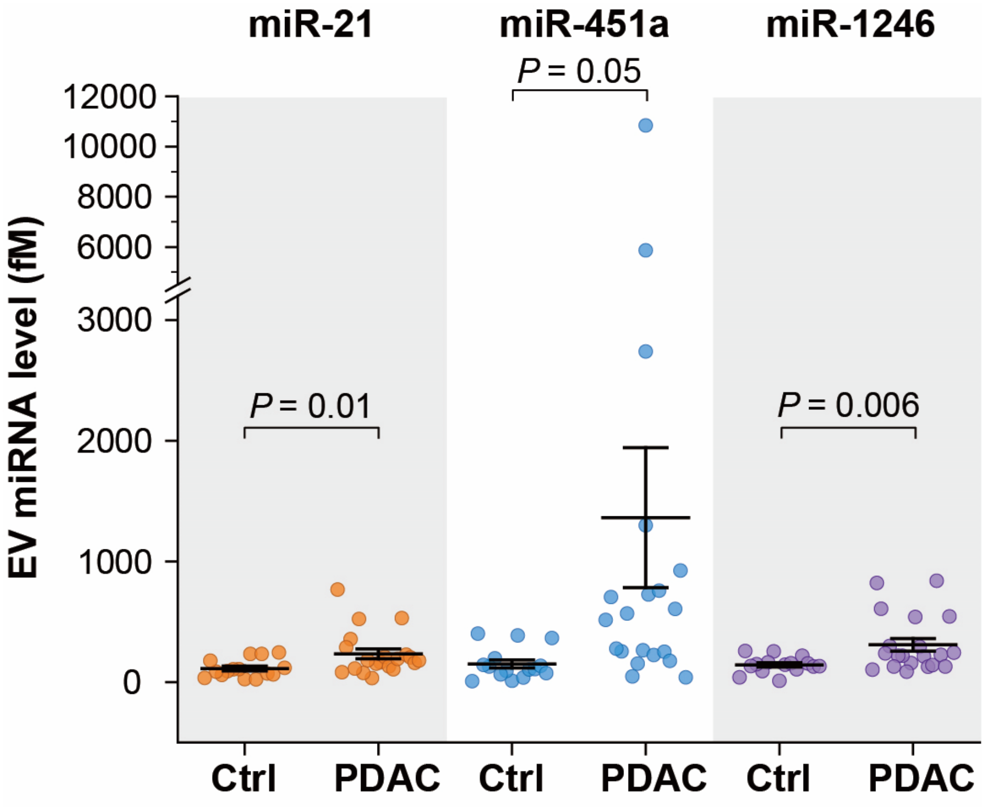


**Figure S9. Scatter plots of the level of individual sEV-miRNAs measured by RT-qPCR.** Each miRNA in each sample was assayed in two technical replicates and the average signals were used to calculate the miRNA concentrations with the calibration curves established in **Figure S7**. The middle line and error bar represent the mean and one s.e.m., respectively. *P* values were calculated by two-tailed Student's *t*-test with Welch correction.

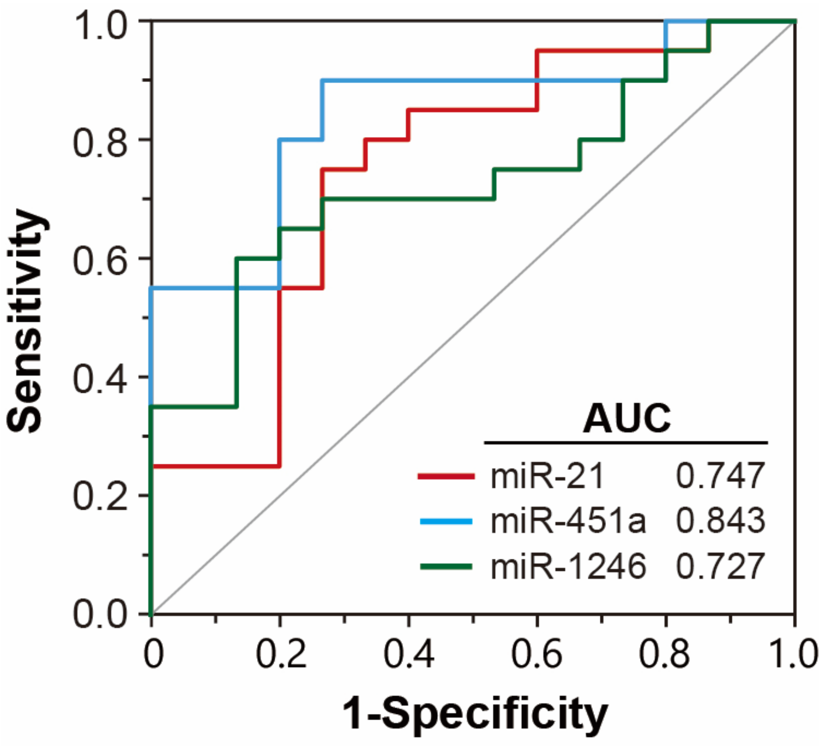


**Figure S10. ROC curves and AUC analysis of individual sEV-miRNAs for PDAC diagnosis.** The ROC curves were plotted using the EV-miRNA levels presented in **Figure S9**. Statistical analyses were performed at 95% confidence level.

**Table S1. Nucleic acid sequences**

| | |
|---|---|
| **DNA Sequences** | |
| Padlock-1 | 5'p-CTGATAAGCTAAGATACCCTAACCATCGATCGTCGCCGTCCAGCTCGACCTCAACATCAGT |
| Padlock-2 | 5'p-CTGATAAGCTAAGATACCCTAACCATCGATTTTACGTCGCCGTCCAGCTCGACCTCAACATCAGT |
| Padlock-3 | 5'p-CTGATAAGCTACGTCGCCGTCCAGCTCGACCAGATACCCTAACCATCGATTCAACATCAGT |
| Padlock-4 | 5'p-CTGATAAGCTAAGATACCCTCGTCGCCGTCCAGCTCGACCAACCATCGATTCAACATCAGT |
| Padlock-miR-196a | 5'p-GAAACTACCTAAGATACCCTAACCATCGATCGTCGCCGTCCAGCTCGACCCCCAACAACAT |
| Padlock-miR-451a | 5'p-GGTAACGGTTTAGATACCCTAACCATCGATCGTCGCCGTCCAGCTCGACCAACTCAGTAAT |
| Padlock-miR-1246 | 5'p-AAATCCATTAGATACCCTAACCATCGATCGTCGCCGTCCAGCTCGACCCCTGCTCCAA |
| Synthetic short cut | GGTCGAGCTGGACGGCGACGATCGATGGTTAGGGTATCTTAGCTTATCAGACTGATGTTGA |
| Reporter | 5-FAM-TTATT-IABkFQ-3' |
| **RNA sequences** | |
| miR-21 | UAGCUUAUCAGACUGAUGUUGA |
| miR-21-mismatch | UAGCUUAUCAUACUGAUGUUGA |
| miR-17 | CAAAGUGCUUACAGUGCAGGUAG |
| miR-18a | UAAGGUGCAUCUAGUGCAGAUAG |
| miR-106a | AAAAGUGCUUACAGUGCAGGUAG |
| miR-145 | GUCCAGUUUUCCCAGGAAUCCCU |
| miR-196a | UAGGUAGUUUCAUGUUGUUGGG |
| miR-205 | UCCUUCAUUCCACCGGAGUCUG |
| miR-451a | AAACCGUUACCAUUACUGAGUU |
| miR-1246 | AAUGGAUUUUUGGAGCAGG |
| crRNA | UAAUUUCUACUAAGUGUAGAUCGUCGCCGUCCAGCUCGACC |

**Table S2. Comparison of EXTRA-CRISPR with other Cas-based miRNA detection methods.**

| Method | Pre-amplification | Cas function | One-pot reaction | # of major components | Assay time | LOD | Cost per test |
|---|---|---|---|---|---|---|---|
| Cas12a-SCR[1] | RCA to generate pre-crRNA | Fluorogenic readout by trans-cleavage | No, 3 steps | 4 enzymes, 6 probes | 6 h | miR-21: 47 fM | $1.9 |
| Cas12a-TCA[2] | HRCA followed by transcription to generate pre-crRNA | Fluorogenic readout by trans-cleavage | No, 3 steps | 4 enzymes, 7 probes | 7 h | miR-21: 1 fM | $2.7 |
| Cas12a-mediated cascade amplification[3] | Ligation-triggered transcriptional amplification and DNAzyme cleavage to generate crRNAs | Fluorogenic readout by trans-cleavage | No, 3 steps | 3 enzymes, 8 probes | 3.5 h | let-7a: 21.9 fM | $3.5 |
| Cas12a-enhanced RCA[4] | Ligation and linear RCA of target | Fluorogenic readout by trans-cleavage | No, 3 steps | 3 enzymes, 3 probes | 4.5 h | miR-21: 34.7 fM | $3.7 |
| RCA-assisted CRISPR/Cas9 cleavage (RACE)[5] | Ligation and linear RCA of target | Fluorogenic readout by Cas9 cleavage | No, 4 steps | 3 enzymes, 3 probes per target | 3.5 h | miR-21: 90 fM | N/A |
| Cas13a-powered electrochemical microfluidic sensor[6] | Amplification-free | trans-cleavage of affinity probe for indirect electro-chemical detection | No, Multiple | 2 enzymes, 2 probes, 1 antibody, 1 sensor chip | 4 h | miR-19b, miR-20a: 10 pM | N/A |
| CRISPR/Cas12a-Assisted Ligation-Initiated LAMP[7] | Ligation and LAMP for exponential target amplification | Fluorogenic readout by trans-cleavage | No, 3 steps | 3 enzymes, 6 probes | 1 h | let-7a: 0.1 fM | $1.0 |
| EXTRA-CRISPR | No | Cis-cleavage of RCA amplicon to create secondary templates; fluorogenic readout by trans-cleavage | Yes | 3 enzymes, 3 probes | 1.5h | miR-21: 1.64 fM; miR-196a: 1.35 fM; miR-451a: 4.14 fM; miR-1246: 7.96 fM | $0.6 |

**Table S3. Summary of the recipes for EXTRA-CRISPR detection of four miRNAs.**

| Components (μL) | miR-21 | miR-196 | miR451a | miR-1246 |
|---|---|---|---|---|
| Buffer (10 ×) | 2 μL | 2 μL | 2 μL | 2 μL |
| dNTP | 400 nM | 400 nM | 400 nM | 400 nM |
| BSA | 0.2 mg/mL | 0.2 mg/mL | 0.2 mg/mL | 0.2 mg/mL |
| Padlock | 100 nM | 100 nM | 10 nM | 10 nM |
| miRNA | 2 μL | 2 μL | 2 μL | 2 μL |
| Phi29 | 0.1 U/μL | 0.1 U/μL | 0.05 U/μL | 0.05 U/μL |
| SplintR | 0.625 U/μL | 1.25 U/μL | 1.25 U/μL | 1.25 U/μL |
| RNP | 1 nM | 1 nM | 1 nM | 1 nM |
| Reporter | 1μM | 2 μM | 4 μM | 1 μM |
| Water | Adjust water volume accordingly (total reaction volume: 20 μL). | | | |

**Table S4. Plasma samples used in this study**

| **Samples for EV miRNA profiling** | | | | | |
|---|---|---|---|---|---|
| **Patients** | **Body site** | **Pathology Status** | **Histologic type** | **Pathological Stage of Cancer** | **Subject Age** |
| 1 | Pancreas, NOS | Primary Cancer | Adenocarcinoma, NOS | pT3N0 | 70 |
| 2 | Pancreas, NOS | Primary Cancer | Adenocarcinoma, NOS | pT4pN1 | 81 |
| 3 | Pancreas, NOS | Primary Cancer | Adenocarcinoma, NOS | pT2pN0 | 75 |
| 4 | Pancreas, NOS | Primary Cancer | Adenocarcinoma, NOS | N/A | 68 |
| 5 | Pancreas, NOS | Primary Cancer | Adenocarcinoma, NOS | N/A | 68 |
| 6 | Pancreas, NOS | Primary Cancer | Adenocarcinoma, NOS | N/A | 54 |
| 7 | Pancreas, NOS | Primary Cancer | Adenocarcinoma, NOS | pT3N1 | 58 |
| 8 | Pancreas, NOS | Primary Cancer | Adenocarcinoma, NOS | pT3N1 | 59 |
| 9 | Pancreas, NOS | Primary Cancer | Adenocarcinoma, NOS | N/A | 73 |
| 10 | Pancreas, NOS | Primary Cancer | Adenocarcinoma, NOS | pT2N0 | 73 |
| 11 | Pancreas, NOS | Primary Cancer | Adenocarcinoma, NOS | pT3pN0 | 55 |
| 12 | Pancreas, NOS | Primary Cancer | Adenocarcinoma, NOS | N/A | 76 |
| 13 | Pancreas, NOS | Primary Cancer | Adenocarcinoma, NOS | N/A | 69 |

| | | | | | |
|---|---|---|---|---|---|
| 14 | Pancreas, NOS | Primary Cancer | Adenocarcinoma, NOS | pT3N0 | 81 |
| 15 | Pancreas, NOS | Primary Cancer | Adenocarcinoma, NOS | pT3 pN1 | 64 |
| 16 | Pancreas, NOS | Primary Cancer | Adenocarcinoma, NOS | N/A | 75 |
| 17 | Pancreas, NOS | Primary Cancer | Adenocarcinoma, NOS | pT3N1 | 65 |
| 18 | Pancreas, NOS | Primary Cancer | Adenocarcinoma, NOS | pT3N1 | 80 |
| 19 | Pancreas, NOS | Metastatic cancer | Adenocarcinoma, NOS | N/A | 40 |
| 20 | Pancreas, NOS | Primary Cancer | Adenocarcinoma, NOS | N/A | 68 |
| **Healthy control** | **Body site** | **Pathology Status** | **Histologic type** | **Pathological Stage of Cancer** | **Subject Age** |
| 21 | N/A | Healthy control | N/A | N/A | 30 |
| 22 | N/A | Healthy control | N/A | N/A | 31 |
| 23 | N/A | Healthy control | N/A | N/A | 27 |
| 24 | N/A | Healthy control | N/A | N/A | 23 |
| 25 | N/A | Healthy control | N/A | N/A | 21 |
| 26 | N/A | Healthy control | N/A | N/A | 42 |
| 27 | N/A | Healthy control | N/A | N/A | 34 |
| 28 | N/A | Healthy control | N/A | N/A | 26 |
| 29 | N/A | Healthy control | N/A | N/A | 33 |
| 30 | N/A | Healthy control | N/A | N/A | 40 |
| 31 | N/A | Healthy control | N/A | N/A | 23 |
| 32 | N/A | Healthy control | N/A | N/A | 36 |
| 33 | N/A | Healthy control | N/A | N/A | 27 |
| 34 | N/A | Healthy control | N/A | N/A | 47 |
| 35 | N/A | Healthy control | N/A | N/A | 46 |

NOS: Not otherwise specified
